# An in vivo screen for proteolytic switch domains that can mediate Notch activation by force

**DOI:** 10.1101/2024.07.10.602225

**Authors:** Frederick C. Baker, Jacob Harman, Trevor Jordan, Breana Walton, Amber Ajamu-Johnson, Rama F. Alashqar, Simran Bhikot, Gary Struhl, Paul D. Langridge

## Abstract

Notch proteins are single pass transmembrane receptors activated by sequential extracellular and intramembrane cleavages that release their cytosolic domains to function as transcription factors in the nucleus. Upon binding, Delta/Serrate/LAG-2 (DSL) ligands activate Notch by exerting a “pulling” force across the intercellular ligand/receptor bridge. This pulling force is generated by Epsin-mediated endocytosis of ligand into the signal-sending cell, and results in the extracellular cleavage of the force-sensing Negative Regulatory Region (NRR) of the receptor by an ADAM10 protease [Kuzbanian (Kuz) in *Drosophila*]. Here, we have used chimeric Notch and DSL proteins to screen for other domains that can function as ligand-dependent proteolytic switches in place of the NRR in the developing *Drosophila* wing. While most of the tested domains are either refractory to cleavage or constitutively cleaved, we identify several that mediate Notch activation in response to ligand. These NRR analogues derive from widely divergent source proteins and have strikingly different predicted structures. Yet, almost all depend on force exerted by Epsin-mediated ligand endocytosis and cleavage catalyzed by Kuz. We posit that the sequence space of protein domains that can serve as force-sensing proteolytic switches in Notch activation is unexpectedly large, a conclusion that has implications for the mechanism of target recognition by Kuz/ADAM10 proteases and is consistent with a more general role for force dependent ADAM10 proteolysis in other cell contact-dependent signaling mechanisms. Our results also validate the screen for increasing the repertoire of proteolytic switches available for synthetic Notch (synNotch) therapies and tissue engineering.

**One sentence summary:** A scalable in vivo screen for Epsin and ADAM10 dependent proteolytic switches that can mediate Notch activation in response to force.

## Introduction

Many processes in biology depend on contact-dependent, juxtacrine signaling in which transmembrane ligands and receptors form intercellular bridges between “sending” and “receiving” cells. Such bridges are likely exposed to pulling forces as one or both ends of the bridge undergo endocytosis and the resulting mechanical tension across the bridge can lead to ectodomain cleavages that regulate signaling. A prototypical example is the receptor Notch, a highly conserved single pass transmembrane protein that has profound and pervasive roles in development, physiology and disease (*1–3*). Notch is activated by single pass transmembrane ligands of the conserved Delta/Serrate/LAG-2 (DSL) family. Upon binding, DSL ligands induce cleavage and shedding of the Notch ectodomain by an ADAM10 protease [Kuzbanian (Kuz) in *Drosophila*] tethered to the surface of the receiving cell (*4–6*). Ectodomain shedding renders the rest of the receptor a substrate for intramembrane cleavage by γ-secretase (*7–9*), releasing the cytosolic domain—a transcription factor—for entry into the nucleus (*10–13*).

For Notch, activation depends on mechanical tension exerted across the intercellular ligand/receptor bridge by endocytosis of the ligand into the sending cell. Specifically, upon binding to Notch, DSL ligands are recruited to the Clathrin endocytic pathway by the adapter protein Epsin (*14–16*). The process of ligand recruitment and/or internalization then generates a pulling force across the ligand/receptor bridge that renders it subject to ADAM10 proteolysis (*17, 18*). Both in vivo and biophysical studies indicate that cleavage depends on the exposure of an otherwise buried site in a conserved, juxtamembrane “Negative Regulatory Region” (NRR) that behaves as a force sensitive proteolytic switch (*17, 19*). Indeed, the key role of the NRR can be recapitulated using a heterologous, force-dependent proteolytic switch—the A2 domain of Von Willebrand Factor (*20, 21*)—in place of the NRR, provided it is tuned to the appropriate force level (*18*).

The ectodomains of many other cell surface proteins are also cleaved by ADAM10 proteases (*22, 23*), raising the possibility that force-dependent, ADAM10 proteolysis plays a more general role in juxtacrine signaling. In support, although many ADAM10 substrate proteins are cleaved constitutively [for example (*24*)], proteolysis of at least some depends on ligand (*5, 25*). However, the sheer number of ADAM10 substrates identified to date [at least 100 (*26*)], as well as the lack of a defined consensus cleavage site (*27*), make it difficult to assess the prevalence of ADAM10-dependent force-sensitive switches in juxtacrine signaling. Moreover, the possibility that any given ADAM10-sensitive domain functions as a proteolytic switch needs to be validated by experiment. Notably, such tests have yielded positive results in a cell culture screen of Sea urchin Enterokinase Agrin-like (SEA) domains, which have structural similarity with Notch NRRs and are found in many transmembrane receptors and other cell surface proteins (*28*). The capacity of SEA domains to function as NRR-like proteolytic switches in cell culture is consistent with a more general role for such domains in force-sensitive signaling events. However, it is unclear how well cell culture conditions capture the native signaling processes in vivo, where the relevant interactions often occur between cells in polarized epithelia, and for which ligand and/or receptor endocytosis may be necessary to generate the requisite, activating force. Also unclear is the structural diversity of domains that act as ADAM10 dependent proteolytic switches.

Here we present the results of an in vivo screen designed to identify protein domains that can function as proteolytic switches in place of the Notch NRR. The screen is predicated on the use of chimeric forms of *Drosophila* Notch and Delta in which the native ligand and receptor ectodomains are replaced by heterologous ligand and receptor binding domains, and productive signaling monitored in vivo by the induction of Notch target genes in wing imaginal discs (*18*). Of 43 chimeric Notch receptors that contain different candidate switch domains, we have identified 11 that respond to ligand in an Epsin and Kuz dependent manner. Strikingly, these proteolytic switches derive from a wide range of proteins and appear remarkably diverse in structure, indicating that the repertoire of possible switch domains is surprisingly large and suggesting that many different cell surface proteins have the potential to be force-dependent targets for Kuz/ADAM10 proteases. Our results also validate the screen as a means to identify new switch domains that can be used to diversify the collection of available synthetic Notch (synNotch) receptors for biomedical applications (*29, 30*).

## Results

### In vivo screen for putative force sensitive proteolytic switches able to substitute for the Notch Negative Regulatory Region (NRR)

To screen protein sequences for their capacity to function as NRR-like proteolytic switches, we generated transgenes that encode chimeric Notch receptors in which (i) the ligand-binding, EGF-repeat domain of the native receptor is replaced by the ligand-binding domain of the <u>F</u>ollicle <u>S</u>timulating <u>H</u>ormone <u>R</u>eceptor (FSHR), and (ii) the juxtamembrane NRR is replaced by candidate switch domains from other proteins (Fig. 1A, B). All such FSHR-Notch receptors retain the transmembrane and intracellular domains of the native Notch receptor (Nicd). Hence, when activated, they drive well-characterized transcriptional and morphological outputs of the Notch signaling pathway. We then asked whether each of these receptors can be activated in vivo by a chimeric form of Delta (Dl) in which the ectodomain is replaced b<u>y</u> Follicl<u>e</u> Stimulating <u>H</u>ormone (FSH-Dl). Importantly, native FSH is a heterodimer of distinct α and β subunits. As in our prior studies (*18, 31*), we generate the functional FSH-Dl ligand by co-expressing an otherwise inactive FSHβ-Dl ligand together with secreted FSHα.

**Figure 1.**
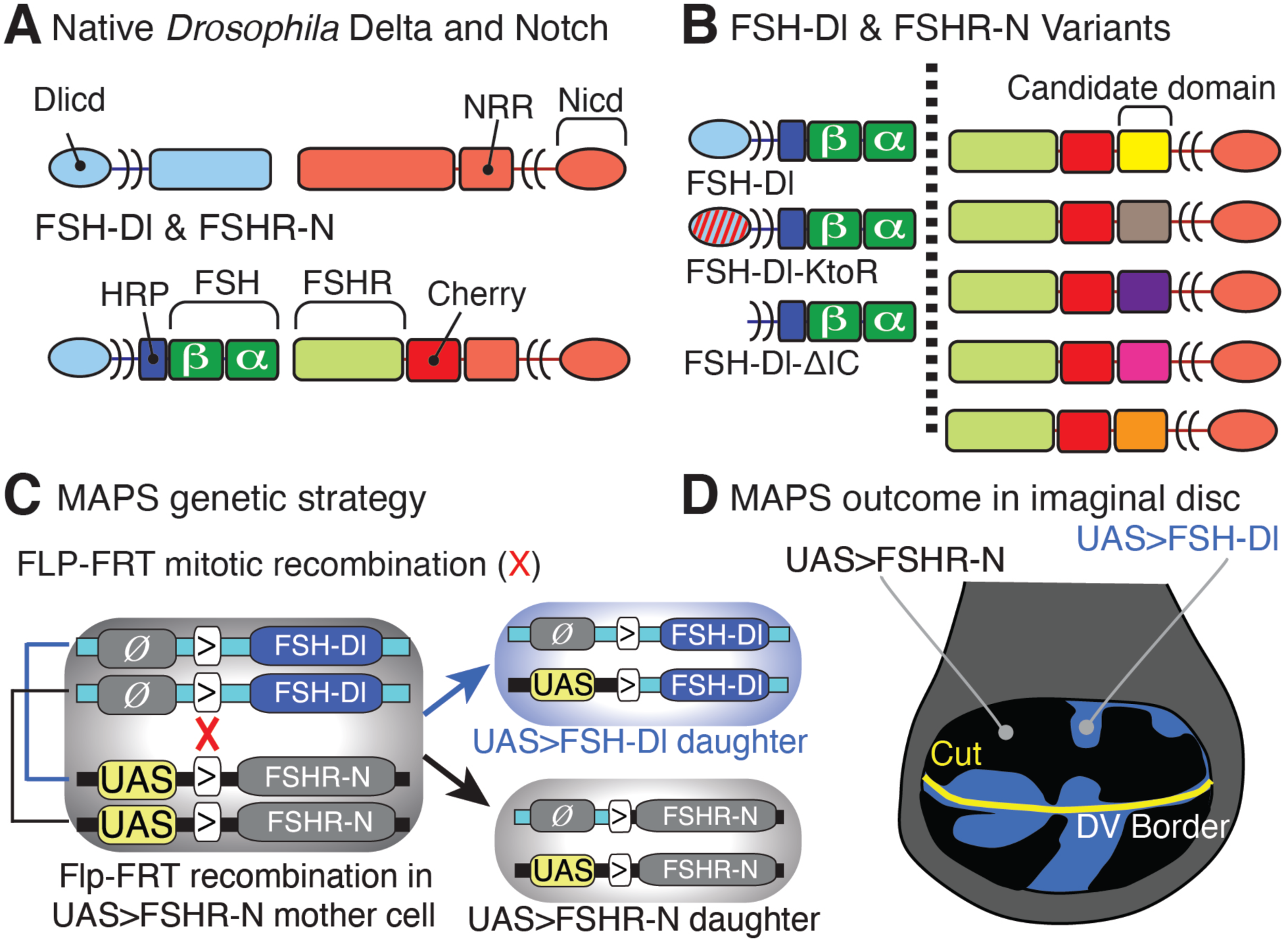
Functional screen to identify force-sensitive cleavage domains in vivo. A. Cartoons of *Drosophila* Delta (Dl)/Notch (N) and FSH-Dl/FSHR-Notch chimeras in which the ligand binding regions of Delta/N were replaced with functional heterologous binding regions. The Dl extracellular domain was replaced by the β-subunit of Follicle Stimulating Hormone (FSHβ) and coexpression of FSHα reconstitutes the composite FSH ligand (FSH). Reciprocally, the ligand-binding (EGF) portion of the N ectodomain was replaced by the FSH receptor ectodomain (FSHR). The intracellular domain of both Notch (Nicd) and Delta (Dlicd) were not altered. B. The canonical FSHR-Notch receptor was modified by replacing the NRR with a single candidate sequence to produce 43 different FSHR-Notch receptors each with distinct juxtamembrane regions. For the screen FSH-Dl was used, but for later analyses FSH-Dl was modified by replacing Dlicd that targets FSH-Dl into the Epsin route with domains that exclude the ligand from that route, either through mutation of the lysines to arginine (FSH-Dl-KtoR) or removal of the intracellular domain entirely (FSH-Dl-ΔIC). The coding sequence for each receptor was placed downstream of a UAS promoter followed by an FRT (>) recombination target sequence to form the *UAS>FSHR-Notch* transgenes, each of which was part of a vector with one or two of several eye (w+, Rap rough eye phenotype), wing (y+), bristle (y+) or cuticle color (y+) marker genes. Mixtures of *UAS>FSHR-Notch* trans genes with distinct markers were introduced at identical genomic docking sites (86Fb) by a single micro-injection. The FSH-Dl coding sequence was placed downstream of an FRT, but no promoter (Ø), to form *Ø>FSHβ-Dl*, and inserted at the 86Fb genomic docking site. C. Mosaic Analysis by Promoter Swap (MAPS) allows the generation of interfaces between clones of FSH-Dl expressing cells and surrounding FSHR-Notch expressing tissue. Specifically, the coding sequences of the chimeric FSHR-Notch receptors were placed downstream of an *Upstream Activating Sequence* (*UAS*) promoter controlled by the yeast Gal4 transcription factor, but separated from the promoter by a target for mitotic recombination catalyzed by the yeast Flp recombinase [FRT, symbolized by “>”]. All of the resulting *UAS>FSHR-Notch* transgenes were inserted at the same attP genomic docking site [at cytological position 86Fb] and in the same orientation as a *Ø>FSH-Dl* transgene that lacks a functional promoter (“*Ø*”). Consequently, transheterozygous *UAS>FSHR-Notch/Ø>FSH-Dl* cells express only FSHR-Notch in response to Gal4. However, when subjected to heat shock induced Flp recombinase, these cells can undergo mitotic recombination across the FRTs (red cross) to generate single cells in which the *UAS* promoter now drives expression of the ligand rather than the receptor. Only one of the two possible segregation outcomes is shown (left brackets); the alternative segregation event also results in an FSH-Dl expressing daughter cell adjacent to sibling and parental cells that express only FSHR-Notch. D. MAPS generated *UAS>FSH-Dl* founder cells develop to form clones in a surround of *UAS>FSHR-Notch* tissue. The prospective wing (blue/black) is divided into cells expressing either FSH-Dl (the HRP epitope tag of FSH-Dl is stained blue) or receptor under Gal4/UAS control. Normally, peak levels of native DSL/N signaling induce expression of the N target gene *cut* along the boundary between the dorsal (D) and ventral (V) compartments (Cut protein, yellow).

Because virtually all of our experiments are performed in the presence of co-expressed FSHα, we refer to the FSHβ-Dl coding sequence and encoded protein—for convenience —as FSH-Dl, except when we perform negative controls in which it is expressed in the absence of FSHα.

To execute this approach at scale, we incorporated three strategies to optimize generating chimeric FSHR-Notch receptors and testing their capacity to respond to FSH-Dl.

First, the coding sequences for a collection of FSHR-Notch chimeric receptors, each having a different protein domain in place of the native NRR, were introduced into a series of transformation vectors bar-coded by using different transgenic markers (for example variously deleted forms of a genomic *yellow^+^* rescuing fragment that generate different patterns of body and bristle pigmentation). This allowed us to micro-inject mixtures of several transgenes at a time, and then distinguish transformants of each transgene by phenotype, thus saving time, materials and cost.

Second, we used transformation vectors designed to be compatible with our Mosaic Analysis by Promoter Swap (MAPS) technique (*18*). This allows us to generate potential signaling interfaces between clones of FSH-Dl expressing cells and surrounding FSHR-Notch expressing tissue generated by Flp-mediated mitotic recombination [described in detail in Fig. 1C; (*18*)].

Third, we used a *nubbin.Gal4* (*nub.Gal4*) transgene to drive expression of *UAS>FSH-Dl* and *UAS>FSHR-Notch* transgenes in the “pouch” of the wing imaginal disc—a circular domain within the disc destined to form the adult wing [Fig. 1D; “>” denotes Flp Recombinase Targets (FRTs)]. Normally, peak DSL/Notch signaling in the pouch is restricted to a thin stripe of cells flanking the dorso-ventral (DV) boundary, where it (i) induces expression of the transcription factor Cut, (ii) specifies formation of wing margin bristles, and (iii) directs local production of Wingless (Wg), a morphogen that controls wing growth (*32–34*). Early heat shock induction of *UAS>FSH-Dl* clones in *UAS>FSHR-Notch* wing discs generates clonal interfaces between dedicated FSH-Dl and FSHR-Notch expressing cells. If a given FSHR-Notch receptor can be activated by FSH-Dl, these interfaces will induce ectopic expression of Notch target genes and altered wing morphology, both of which are easy to score in vivo.

### Protein domains tested for NRR-like proteolytic switch properties

We chose to test domains from a diverse collection of proteins to assess the efficacy of the screen as well as the repertoire of structures that can mediate Notch cleavage in response to force (Table 1). Kuzbanian/ADAM10 proteolysis typically targets the juxtamembrane extracellular regions of cell surface proteins, and the amino acid sequence cleaved varies among substrates. Hence, we mostly tested domains positioned immediately N-terminal to the transmembrane domain of cell surface proteins involved in juxtacrine signaling and did not restrict the choice to domains with particular structural features. We also included a number of miscellaneous peptide sequences and protein domains. Candidate domains differed in length and structural composition, ranging from juxtamembrane regions from surface receptors (for example from Eph receptors) to proteolytic switches previously identified in cell culture [table S1, (*28*)], to unstructured domains (for example flagelliform, a spider silk protein that is a concatemer of glycine-proline-alanine repeats), to protein domains suggested by serendipitous findings (for example Venus fluorescent protein), to a few truncated versions of the native Notch NRR. Finally, for benchmarks, we used the intact NRRs of *Drosophila* Notch and the *C.elegans* Notch homologue LIN-12, which we have characterized in previous work (*18, 31*). Both are strictly ligand-dependent, but the Notch NRR is tuned to a force threshold requiring the Epsin pull, whereas the LIN-12 NRR is tuned to a lower threshold that does not. We present, below, the results of a screen of 43 *UAS>FSHR-Notch* transgenes, each encoding a different candidate proteolytic switch domain.

**Table 1.** List of domains in the ligand-dependent and constitutive categories. Columns describe the domain’s abbreviated name, the identity of the native source protein and species, the size of the domain (#AA=number of amino acids), a brief description of the native source protein and the reason for its inclusion in the screen. Candidate domains cleaved in response to ligand are shown in green, including domains in Class I (high force tuning, dark green), II (low force tuning, light green) and III (‘hyperactive’, chartreuse). Note that the FSHR-Notch receptors carrying the candidate domains have a Cherry tag that showed these receptors had similar subcellular distributions and levels of accumulation, with any modest differences not correlating with their assignment into one of three distinct classes of response to ligand. Candidate domains cleaved constitutively are indicated in yellow. Domains that were not cleaved (see also table S2) include FSHR-Notch lacking the LNR, EprinA2, *Drosophila* Fat, Filamin, Neuregulin, Starry Night, Dachsous, junctional adhesion molecule A, prion related protein, human DSCAM, viral spike proteins, epithelial cellular adhesion molecule, KIAA0319, and Muc13. See also table S1 for comparison to the SNAPs cell culture assay (*28*).

| Name | Domain Source | Species | Size (#AA) | Description | Reason for inclusion in screen |
| --- | --- | --- | --- | --- | --- |
| DCC | Deleted in Colorectal Cancer/ Frazzled | *Drosophila* | 18 | Neuronal guidance receptor | Evidence that ICD acts as a transcription factor |
| EphB2 | Ephrin-type receptor B2 | Mouse | 110 | Neuronal guidance receptor | Native receptor is cleaved in response to ligand binding |
| DEph | Ephrin-type receptor | *Drosophila* | 250 | Neuronal guidance receptor | Native receptor is cleaved in response to ligand binding |
| Serrate | Serrate | *Drosophila* | 264 | Notch ligand | Cleaved by Kuz |
| FAT1 | FAT1 | Human | 93 | Atypical cadherin | SNAPS: Cleavage in response to ligand |
| Dscam1 | Down Syndrome Cell Adhesion Molecule 1 | *Drosophila* | 220 | Neuronal guidance receptor | Evidence that ICD acts as a transcription factor |
| Venus GFP | Venus GFP | *Aequorea victoria* | 242 | Entire coding region of Venus GFP | Serendipitous result |
| DAG | Dystroglycan | Human | 261 | Cell adhesion and ECM binding protein | SNAPS: Cleavage in response to ligand |
| NRR<sup>ΔLNR</sup> | Notch variant | *Drosophila* | 45 | Variant of the NRR of Notch | LNRs removed, but S2 site remains |
| ephrinB | Ephrin B1 ligand | Human | 74 | Neuronal guidance protein | Native ligand is cleaved in response to ligand binding |
| Flagelliform | Flagelliform F40 | Spider | 44 | Spider silk – 40 PGGAG repeats | Biophysically demonstrated to stretch at pN forces |
| Dephrin | Ephrin B ligand | *Drosophila* | 211 | Neuronal guidance protein | Native ligand is cleaved in response to ligand binding |
| Robo | Roundabout 1 | *Drosophila* | 371 | Neuronal guidance receptor | DCC/Frazzled related protein |
| Nrg1 | Neuregulin (Nrg1 type III) | Human | 101 | Neuronal guidance protein | ADAM10 substrate |
| Neogenin | Neogenin | Mouse | 367 | Neuronal guidance receptor | ADAM10 substrate |
| Ephrin B2L | Ephrin B2 (Long) | Mouse | 342 | Neuronal guidance protein, juxtamembrane region and 2 fibronectin domains | Cleavage in response to receptor binding |
| Ephrin B2S | Ephrin B2 (Short) | Mouse | 110 | Neuronal guidance protein, juxtamembrane region | Cleavage in response to receptor binding |
| EphB1 | Ephrin B1 receptor | Human | 124 | Neuronal guidance receptor | Receptor cleavage in response to ligand binding |
| Delta | Delta | *Drosophila* | 34 | Notch ligand | Cleaved by Kuz |
| APP | Amyloid-beta precursor protein | Human | 240 | Neuronal repair | ADAM10 substrate |
| CD16 | CD16 | Human | 194 | Immune cell receptor | ADAM10 substrate |
| CD44 | CD44 | Mouse | 461 | Cell adhesion glycoprotein | ADAM10 substrate |
| Cdhr2 | Cadherin-related family member 2 | Human | 356 | Cell adhesion protein | SNAPS: Cleavage in response to ligand |

### Candidate proteolytic switches identified based on adult wing phenotype and Notch target gene expression

To test the capacity of the different FSHR-Notch chimeric receptors to respond to FSH-Dl, females carrying each transgene were outcrossed to “tester” males carrying the *nub.Gal4*, *UAS.FSHα* and *hsp70.flp* transgenes, as well as a *Ø>FSH-Dl* transgene, which encodes FSH-Dl, but lacks a functional promoter (“*Ø*”). Their progeny were then heat-shocked early in larval life to induce expression of the *hsp70.flp* transgene and generate *UAS>FSH-Dl* clones in *UAS>FSHR-Notch* wing discs, and the resulting progeny were screened for abnormal wing phenotypes indicating ectopic FSHR-Notch activity (Fig. 2A).

**Figure 2.**
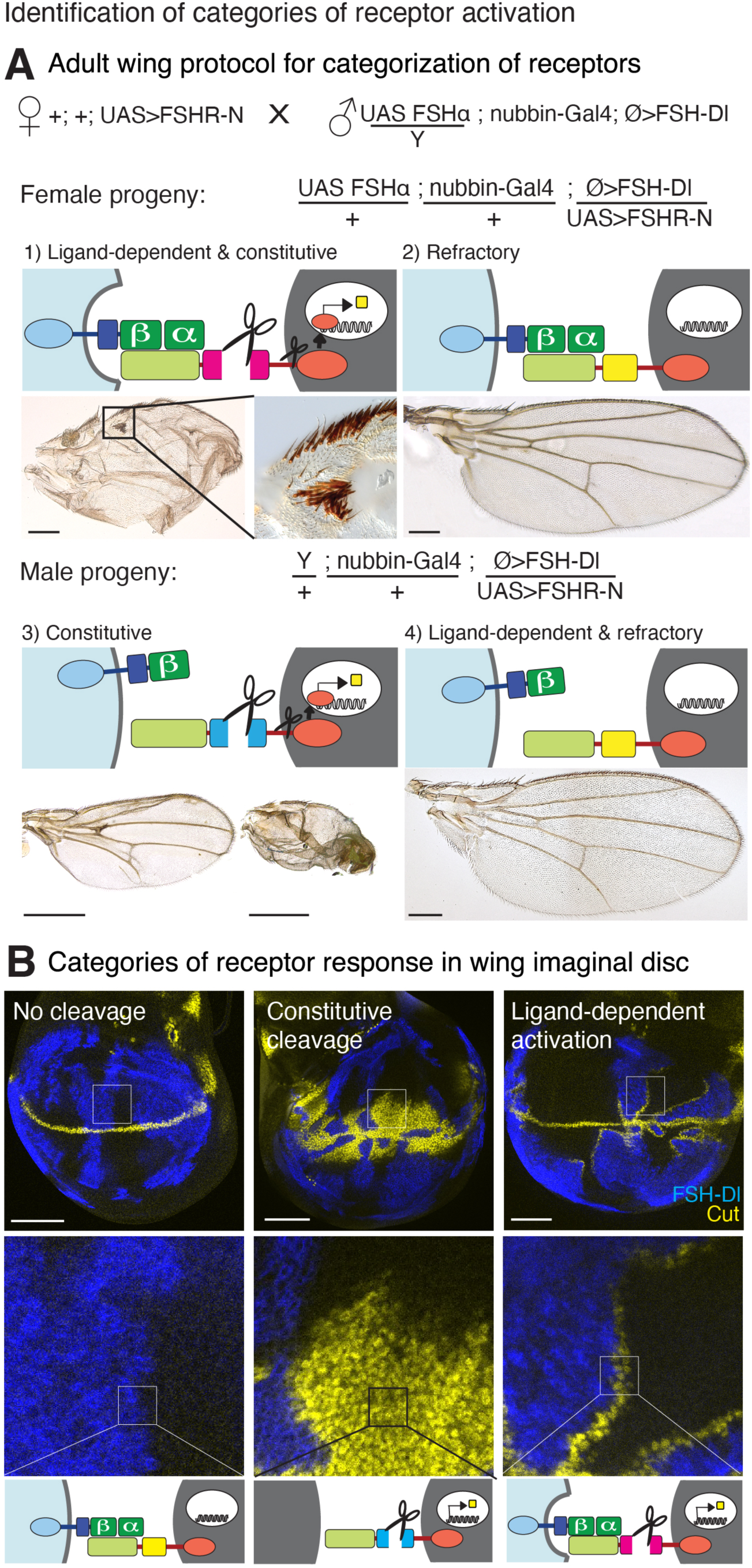
Identification of categories of receptor activation. **A**. To screen for receptor activation in adults, G1 female flies were crossed with male flies from a ligand tester stock with the components for both UAS/Gal4 expression in the wing and the FSH-Dl ligand: *UAS FSHα* on the X chromosome, the nubbin-Gal4 driver on the second chromosome, and the *Ø>FSHβ-Dl* transgene. FSH-Dl/FSHR-Notch interfaces were induced by MAPS and receptor activation categorized by examining adult wing phenotypes resulting from the ectopic activation of Notch target genes: 1) Ligand-dependent and constitutive FSHR-Notch activation in the female progeny leads to ectopic wing margin and smaller wings. 2) An FSHR-Notch receptor that is refractory to cleavage when confronted with ligand produces wildtype wings in the female offspring. 3) Male progeny do not inherit the X chromosome carrying the *UAS FSHα* transgene and do not express functional FSH-Dl. If the FSHR-Notch receptor is constitutively active, receptor activation can occur even in the absence of the *UAS FSHα* transgene and male flies in this case have an abnormal wing phenotype, although to different degrees according to the extent of activation and the random position of the receptor clones. Examples of adult wings with low and high activity levels are shown, with the major phenotype being a reduction in size, change in wing shape and ectopic wing margin. 4) In the absence of the *UAS FSHα* transgene, FSHR-Notch receptors that are ligand-dependent are not activated, resulting in a wildtype wing. Receptors refractory to cleavage will also produce wildtype wings. Scale bar 200 μm. **B.** Patterns of receptor activation in the wing imaginal disc were divided into one of three main categories. Left column, no cleavage of the candidate domain. Clones of FSH-Dl do not produce ectopic Cut at the interface with FSHR-Notch cells. Native Cut remains at the DV boundary. Center column, constitutive cleavage allows FSHR-Notch activation regardless of the presence of ligand, resulting in ectopic Cut produced in all FSHR-Notch cells whether they are close to FSH-Dl cells or not. Right, ligand-dependent activation induces cleavage of FSHR-Notch and ectopic Cut is produced only in FSHR-Notch cells adjacent to FSH-Dl cells. Scale bar 50μm.

*UAS>FSHR-Notch* transgenes that score positively in the wing assay encode FSHR-Notch receptors that are either ligand-dependent or constitutively active. These two possibilities were distinguished in the test cross of *UAS>FSHR-Notch* females to *UAS.FSHα; Ø>FSH-Dl* males as only female progeny co-express FSHα, which is required for ligand activity (Fig. 2A, top), whereas male progeny do not. Accordingly, *UAS>FSHR-Notch* transgenes that cause abnormal wings in both male and female progeny encode constitutively active receptors, whereas those that do so only in female progeny encode ligand-dependent receptors (Fig. 2A, bottom).

To corroborate and extend our classification of FSHR-Notch receptors that scored positively based on the initial wing assay, we repeated the MAPS experiments, this time analyzing the expression of the Notch target gene, *cut*, as monitored by staining for Cut protein, in the developing wing imaginal disc. Cut expression in the wing primordium is normally restricted to a thin strip of cells along the dorso-ventral (D/V) compartment boundary in response to peak Notch activation (Fig. 1D). Hence, both ligand-dependent and constitutively active FSHR-Notch receptors should cause ectopic Cut expression, whereas receptors that are refractory to activation should not. By design, the canonical FSH-Dl and FSHR-Notch proteins are epitope tagged, respectively, with Horse Radish Peroxidase (HRP) and Cherry fluorescent protein (Fig. 1A). As a consequence, both *UAS>FSH-Dl* clones and the surrounding *UAS>FSHR-Notch* tissue can be identified with single cell resolution by monitoring HRP and/or Cherry expression, allowing inductive signaling and constitutive activity to be assayed by monitoring for ectopic Cut expression at or away from the clone borders.

Of the 43 *UAS>FSHR-Notch* transgenes tested, most (40/43) fell into one of three distinct categories based on both Cut expression in the wing imaginal disc (Fig 2B) as well as the adult wing phenotype (Tables 1 and S2): (i) 8/43 showed ectopic Cut restricted to FSHR-Notch expressing cells located near or next to FSH-Dl expressing cells indicating ligand-dependent activation, (ii) 12/43 showed ectopic Cut in FSHR-Notch expressing cells regardless of the presence of FSH-Dl expressing cells indicative of constitutive activation, and (iii) 20/43 receptors showed no ectopic Cut expression indicating the encoded receptors were refractory to activation. However, the remaining 3/43 transgenes were exceptional in encoding receptors that show evidence of both constitutive activity as well as the capacity to respond to ligand, as described below. Hence, they fall into to a special class of “hyperactive” receptors, increasing the number we designate as ligand-dependent from 8/43 to 11/43 (Table 1; chartreuse).

*UAS>FSHR-Notch* transgenes that score negatively encode receptors that are refractory to activation by ligand, consistent with the candidate domain not deriving from a proteolytic switch. However, a bona fide force sensor might nevertheless score negatively for a number of reasons. These include: (i) it’s tuned to a sufficiently high force threshold such that it cannot be activated by FSH-Dl [for example, as is the case for the native A2 domain from VWF (*18*)], (ii) cleavage occurs at a site positioned sufficiently far away from the receptor transmembrane domain to preclude cleavage by γ-secretase, which depends on the size of the remaining ectodomain stub (*9*), or (iii) the candidate domain being tested is inadvertently truncated or impaired by adventitious interactions with neighboring protein sequences to render it inoperable as a switch domain. Hence, we draw no conclusions about the receptors encoded by *UAS>FSHR-Notch* transgenes that tested negatively, but instead, have focused our analysis exclusively on the *UAS>FSHR-Notch* transgenes that yielded positive results.

### Reliance of ligand-dependent FSHR-Notch receptors on Epsin-mediated ligand endocytosis

Given our prior evidence that cleavage of the native NRR depends on the force generated by Epsin-mediated ligand endocytosis (*18*), we next asked if activation of the 11 ligand-dependent FSHR-Notch receptors depends on Epsin. Signaling by DSL ligands normally depends on ubiquitination of cytosolic lysine residues for recruitment to the Clathrin-endocytic pathway by Epsin (*15, 16*). Hence, native DSL ligands in which all lysines in the cytosolic domain are replaced by arginine have greatly reduced capacity to enter the Epsin pathway, whereas those that lack a cytosolic domain are entirely unable to do so (*35, 36*). In the context of FSH-Dl/FSHR-Notch signaling in the wing disc, the Notch and LIN-12 NRR benchmark receptors appear to define opposite ends of an Epsin-dependent force spectrum. Both the FSH-Dl-KtoR and FSH-Dl-ΔIC ligands fail to induce detectable activation of the Notch NRR receptor (*18*), whereas both are similarly effective as the wildtype FSH-Dl ligand in activating the LIN-12 NRR receptor (Langridge et al, 2022). In contrast, FSHR-Notch receptors with modified NRRs that we interpret as having intermediate force thresholds respond moderately to the KtoR ligand and weakly or not at all to the ΔIC ligand. Accordingly, we used the capacity of the 11 ligand-dependent FSHR-Notch receptors to respond wildtype, KtoR, and ΔIC forms of FSH-Dl to assess the force sensitivity of their candidate proteolytic switch domains.

To perform this comparison, we have taken advantage of our previous finding that the distance from the DV boundary that FSH-Dl/FSHR-Notch signaling can induce ectopic Cut expression provides an indication of the strength of signaling (*18*). To normalize for variations in wing disc size, this measurement is expressed as the ratio of the maximum distance of ectopic Cut expression from the DV boundary relative to the length of the DV border. Previously, we used this assay to provide evidence that the benchmark *Drosophila* Notch and LIN-12 NRRs are tuned to different force thresholds [Fig. 3A, (*18, 31*)]. Using this approach, we found that the 11 candidate proteolytic switch domains fell into three distinct classes of force sensitivity.

**Figure 3.**
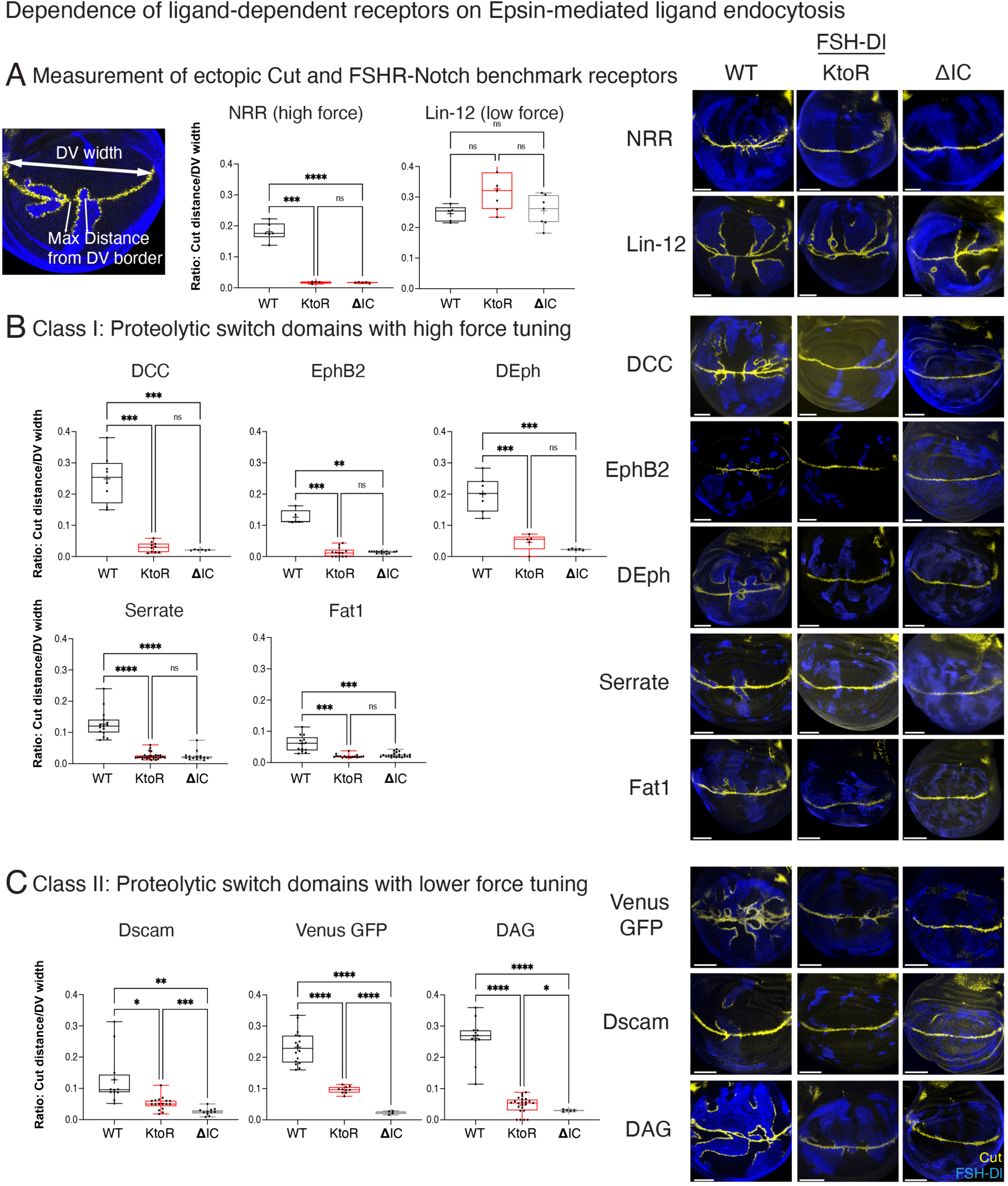
Dependence of ligand-dependent receptors on Epsin-mediated ligand endocytosis. **A**. Quantitation of signaling between FSH-Dl and FSHR-Notch receptors carrying domains cleaved in response to ligand. This was measured as the distance ectopic Cut extends away from the DV boundary relative to disc size (width of Cut expression at the DV boundary) and presented as a ratio of the two distances (left). Signal strength is depicted as boxplots (right) that show the “cut distance/DV width” ratio for each disc as a point, as well as the mean (cross), median (horizontal line), and 25%-75% interquartile ranges (tops and bottoms of the box) for each receptor/ligand combination. Statistical comparisons of the average Cut distance/DV width ratios for each ligand/receptor pair were performed by Brown and Forsythe tests followed by Dunnett TW post-hoc analysis (see methods); *p<0.05, **p<0.01, ***p<0.001, and ****p<0.0001. Each receptor was confronted with three different FSH-Dl variants (Fig. 1A) with intracellular domains corresponding to the wildtype Dl domain (WT), the KtoR mutated Dl domain (KtoR) and a deletion of the intracellular domain (ΔIC). Representative images of these responses are shown on the right. As benchmarks, the response to FSH-Dl ligands of the canonical FSHR-Notch receptor with the *Drosophila* NRR (NRR) and FSHR-Notch with the LIN-12 NRR (LIN-12) is shown. The NRR receptor is strictly Epsin-dependent and does not respond to versions of FSH-Dl that are excluded from the Epsin-route, indicating a high force tuning. In contrast, the LIN-12 responds strongly to all three FSH-Dl ligands, indicating a lower force tuning. **B.** Receptors with a high dependence on Epsin-mediated ligand endocytosis (Class I) have no response to ligand excluded from the Epsin route, similar to the response of the canonical FSHR-Notch with the *Drosophila* NRR. **C.** Receptors with a lower dependence on Epsin-mediated ligand endocytosis (Class II) have small but detectable responses to ligand excluded from the Epsin route. Scale bar 50μm.

Class I receptors (5/11) behaved similarly to the native Notch NRR and contained protein domains from the receptors <u>D</u>eleted in <u>C</u>olorectal <u>C</u>ancer (DCC), Ephrin-type receptor <u>B2</u> (EphB2) and *Drosophila* Eph (DEph), the *Drosophila* DSL ligand Serrate (Ser), and the human protocadherin FAT1. Specifically, they show a readily detectable response to FSH-Dl, but little or no response to FSH-Dl-KtoR or FSH-Dl-ΔIC, indicating a strong, if not absolute requirement for the Epsin-dependent “pull” (Fig. 3B). This is similar to what we have observed previously for both the native Notch NRR as well as disease-associated mutant forms of the A2 domain from Von Willebrand Factor [such as A2^E1638K^, (*18*)], consistent with all 5 of the newly identified class I domains being tuned to respond to force thresholds on par with that of the native Notch NRR.

Class II receptors (3/11) differ from class I receptors in showing a modest response to FSH-Dl-KtoR and weak or no response to FSH-Dl-ΔIC (Fig. 3C). Hence, these switches appear intermediate in their force tuning between the native *Drosophila* Notch NRR, which is strictly Epsin-dependent, and the *C. elegans* Notch NRR LIN-12, which appears Epsin-independent [Fig. 3A, (*31*)]. These class II domains came from Venus fluorescent protein (Venus), Down Syndrome Cell Adhesion Molecule (Dscam) and Dystroglycan.

Class III receptors (3/11) are those we designate as hyperactive, showing both ligand-dependent and constitutive activity. They carry candidate switch domains from Flagelliform, EphrinB and a partial deletion of the fly Not<u>ch</u> NRR <u>(Δ</u>LNR) that retains the native Kuz target site but removes the three conserved Lin-12 Notch Repeats (LNR) that normally occlude it. For these receptors, we find that Cut is present at peak levels in FSHR-Notch expressing cells near FSH-Dl or FSH-Dl KtoR expressing cells but at lower levels further away (Fig. 4A). In the absence of ligand, ectopic Cut expression can be detected, but it is restricted to the vicinity of DV boundary, where we infer low level constitutive activity of the FSHR-Notch receptor boosts the native response of endogenous Notch to Dl above a threshold necessary to generate detectable Cut expression (Fig. 4B). Hence, we interpret these class III hyperactive receptors as having low constitutive activity as well as a readily detectable response to ligand, and this forms a sub-category of the ligand-dependent receptors (Table 1, chartreuse).

**Figure 4.**
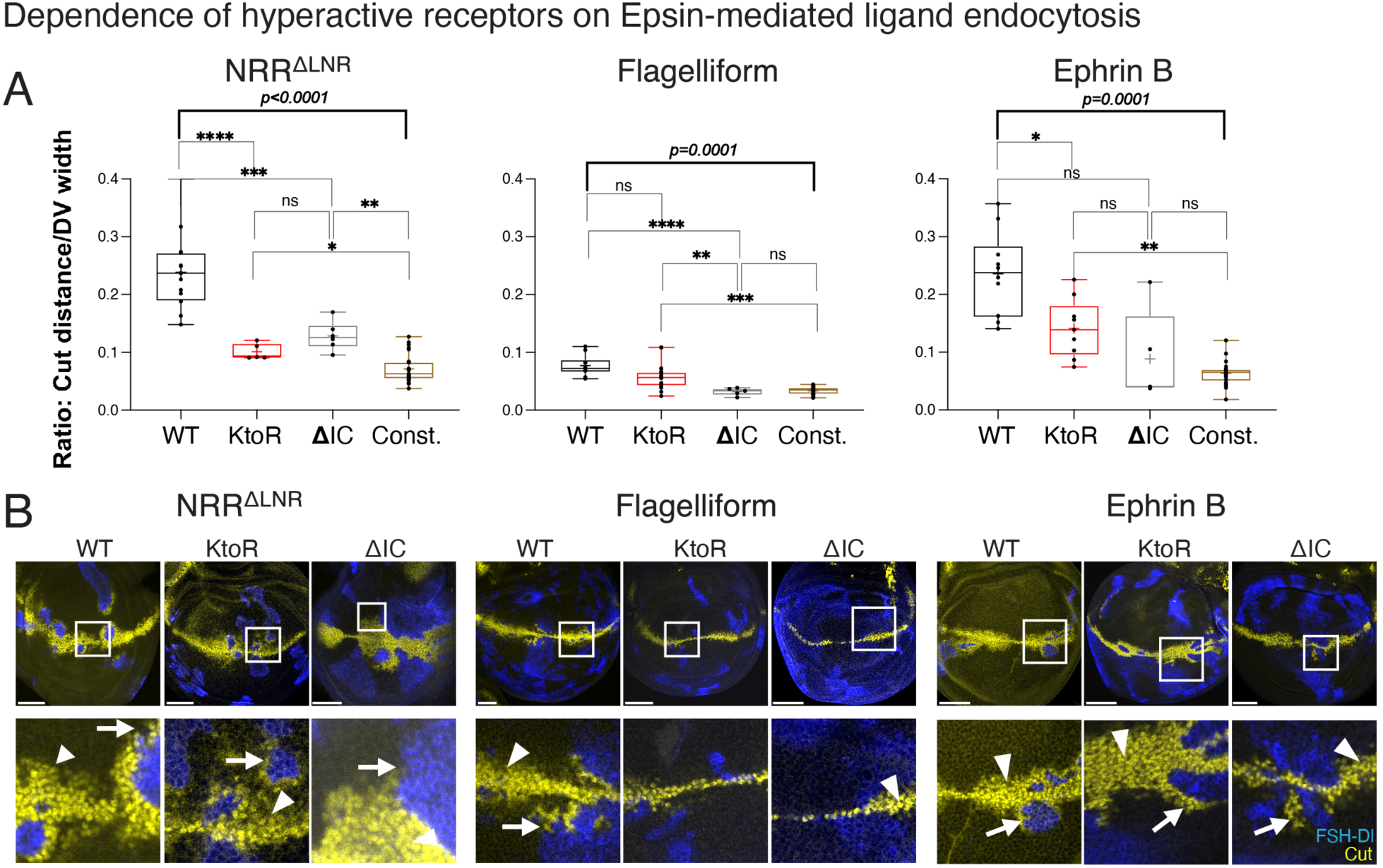
Dependence of hyperactive receptors on Epsin-mediated ligand endocytosis. **A**. Quantitation of signaling between FSH-Dl and FSHR-Notch ‘hyperactive’ variants as in Fig. 3A. Each receptor was confronted with three different FSH-Dl variants with intracellular domains corresponding to the wildtype Dl domain (WT), the KtoR mutated Dl domain (KtoR) and a deletion of the intracellular domain (ΔIC). In addition, the level of constitutive activation of each receptor (Const.) was measured as the maximum distance of ectopic Cut from the DV boundary in regions of the wing away from clonal interfaces. Statistical comparisons of the average Cut distance/DV width ratios for each ligand/receptor pair were performed by Brown and Forsythe tests followed by Dunnett TW post-hoc analysis (see methods); *p<0.05, **p<0.01, ***p<0.001, and ****p<0.0001. **B.** Representative images showing Cut expression in MAPS mosaic wing discs for hyperactive FSHR-Notch receptors paired with FSH-Dl, FSH-Dl-KtoR or FSH-Dl-ΔIC. Significant ectopic Cut is detectable in cells away from FSH-Dl cells (arrowhead) and the maximum distance this occurs is presented in A as the extent of constitutive receptor response (Const.). Ectopic Cut extends furthest from the DV boundary where the receptor cells meet ligand cells (arrow), although the capacity of ligands excluded from the Epsin route (K>R or ΔIC) to elicit a response differs for each receptor. Scale bar 50μm.

We note that FSHR-Notch receptors carrying different ligand-dependent switch domains show a wide range of responses to FSH-Dl, as monitored by the distance over which Cut can be induced away from the DV boundary. However, we don’t know where the different candidate domains are cleaved, which determines the susceptibility of the truncated receptor to Presenilin-dependent transmembrane cleavage (*9*). Further, the expression level or surface distribution of different receptors may vary. Therefore, we do not draw any inferences about differences in force tuning based on comparing the spatial responses of different FSHR-Notch receptors to FSH-Dl. Instead, we draw such inferences only from comparing the responses of the same FSHR-Notch receptor to FSH-Dl versus FSH-Dl-KtoR or FSH-Dl-ΔIC, which we interpret as indicating the relative reliance on Epsin-dependent endocytic force.

### Candidate force sensitive domains require the ADAM protease Kuzbanian for cleavage

The 11 force sensitive candidate domains, as well as the 12 domains that are constitutively cleaved, have little common sequence similarity (Fig. S1) and remarkable structural diversity based on AlphaFold predictions (Fig. S2). This could be because there are few, if any, domain-intrinsic sequence and structural prerequisites for cleavage by ADAM10/Kuz, or because the candidate switch domains are cleaved by other proteases. To resolve this uncertainty, we tested if FSHR-Notch activity mediated by these domains depends on Kuz.

To do this we employed permutations of MAPS technology, one for the ligand-dependent receptors and another for the constitutively active receptors. For both, we depended on the following RNA knock-down approach to greatly reduce Kuz activity in FSHR-Notch expressing cells.

Briefly, expression of *UAS.Kuz-RNAi* under *nub.Gal4* control in otherwise wildtype wing discs renders Cut undetectable at the DV boundary, indicating strongly reduced Kuz activity. However, this knock-down can be rescued cell-autonomously by co-expression of a *UAS.KuzHA* transgene, which is expressed at a sufficiently high level to escape *UAS.Kuz-RNAi* knock-down of both endogenous *Kuz* and *UAS.KuzHA* transcripts (*37*). Hence, in animals that are otherwise wildtype for the native *Kuz* gene, *UAS.KuzHA/UAS.Kuz-RNAi* transheterozygous cells are wildtype for Kuz function and express HA tagged Kuz, whereas homozygous *UAS.Kuz-RNAi/UAS.Kuz-RNAi* cells do not express HA tagged Kuz and have greatly diminished endogenous Kuz function.

Thus, by generating transheterozygotes in which *UAS.Kuz-RNAi* is distal to *UAS>FSHR-Notch* on one chromosome and *UAS.KuzHA* is distal to *Ø>FSH-Dl* on the other, we were able to compare the ability of FSH-Dl expressing cells to induce Cut expression in abutting FSHR-Notch expressing cells depending on whether they retain or lacked Kuz activity (Fig. 5A; described in detail in Fig. S3).

We applied this test and found that all 8 strictly ligand-dependent FSHR-Notch receptors as well as the benchmark receptors containing the NRRs from *Drosophila* Notch and *C. elegans* LIN-12, could activate Cut expression in response to FSH-Dl from abutting signal-sending cells, but only if they retained Kuz function (Fig. 5B). Hence, ligand-dependent activation of all 8 receptors, as well as the two benchmark receptors, depends on Kuz.

We also examined the Kuz requirement for activation for all three hyperactive receptors and a selection of 6 constitutively cleaved receptors (Fig. 6A-D). For the three hyperactive receptors, the position of the receptor clone relative to the ligand clone is not crucial since each receptor shows modest activity in the absence of ligand. We therefore compared activation of the receptors in the presence or absence of the *UAS.Kuz-RNAi* transgene, using a “twin-spot” MAPS strategy that allowed us to assess both the weak constitutive and strong ligand-dependent responses (Fig. 6A). All three hyperactive receptors behaved as strictly Kuz dependent in this assay, as was also the case for control receptors carrying either the native Notch NRR or the E1638K mutant form of the VWF A2 domain (Fig 6B).

**Figure 5.**
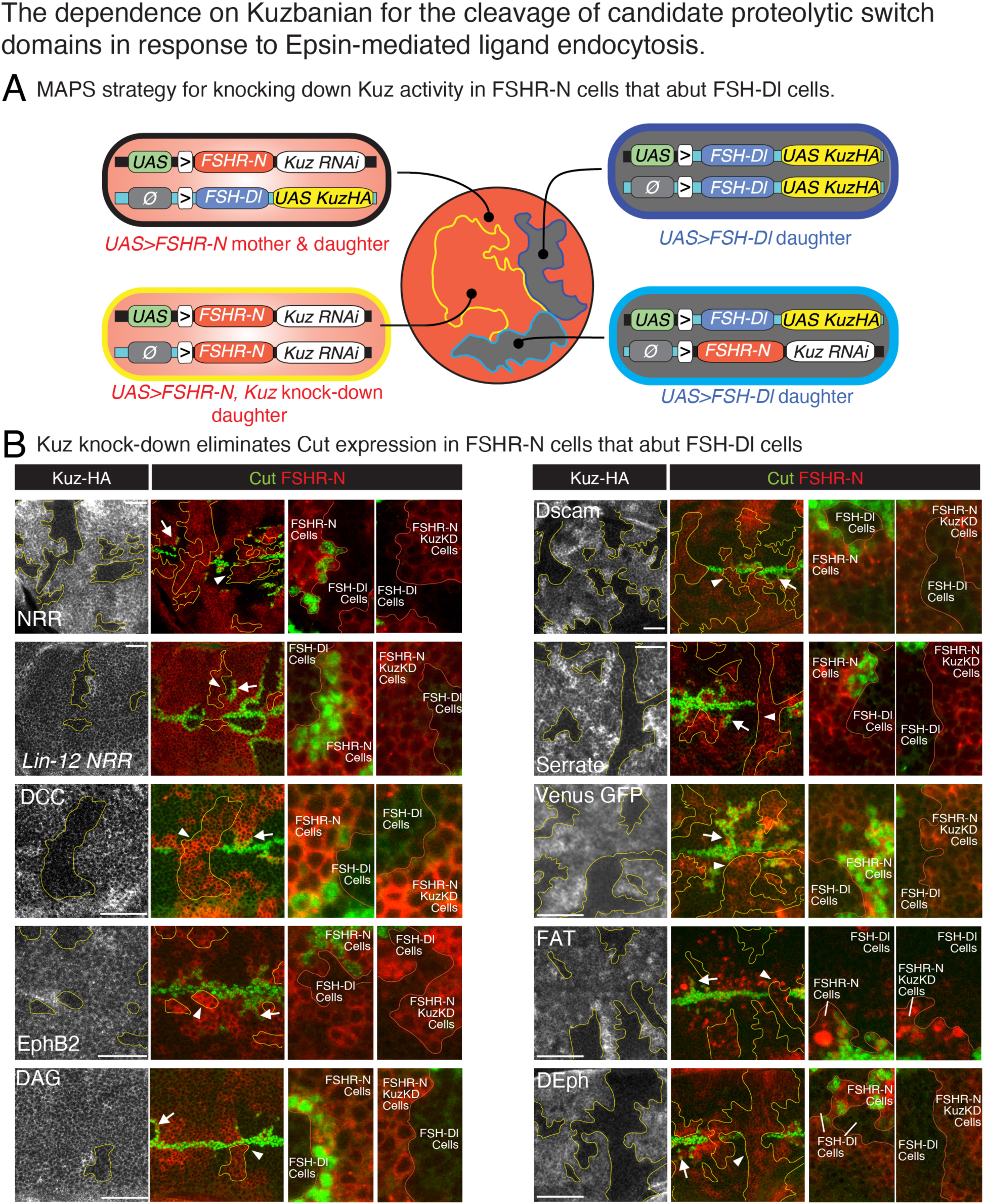
The dependence on Kuzbanian for the cleavage of candidate proteolytic switch domains in response to Epsin-mediated ligand endocytosis. **A**. MAPS technology used to knock-down Kuz activity in genetically marked clones of FSHR-Notch expressing cells. The expression of *UAS.Kuz-RNAi* (Kuz-RNAi) in the wing disc reduces native Notch activation sufficiently to abolish endogenous Cut at the DV boundary, whilst co-expression of *UAS.KuzHA* rescues native DSL/Notch signaling. In background, transheterozygous “mother” cells, one chromosome carries *UAS>FSHR-Notch* with *UAS.Kuz-RNAi* positioned distally and the other chromosome carries *Ø>FSHβ-Dl* with *UAS.KuzHA* positioned distally (upper left). Cells of this “background” genotype express FSHR-Notch but not FSH-Dl and retain KuzHA function. Mitotic recombination across the FRTs generates four kinds of daughter cells, depending on the two possible segregation events (Fig S3). Two of the four, outlined in blue, are “signal-sending” cells that express FSH-Dl but not FSHR-Notch and retain Kuz function. The remaining two are “signal-receiving” cells that express FSHR-Notch but not FSH-Dl, one of which is genetically identical to the mother cell and hence retains KuzHA function (outlined in black), whereas the other is homozygous for *UAS.Kuz-RNAi* and subject to Kuz RNAi knock-down (outlined in yellow). **B**. Comparison of the signal-receiving cells (marked by Cherry, red) in which Kuz is either retained (marked by HA, white) or knocked-down (marked “black” left column by the absence of HA). For each of the FSHR-Notch receptors shown, only clones that retain Kuz activity can express Cut (green) in response to abutting FSH-Dl signal-sending cells (marked “black” by the absence of Cherry). Arrows indicate the Cut response in Kuz positive receiving cells and arrowheads indicate the lack of response in neighboring Kuz knock-down receiving cells (shown at higher magnification to the right, with the mosaic border indicated in red). Note that where Kuz knock-down cells cross the DV boundary they abolish the normal Cut response as expected. The identities of the candidate proteolytic domains are indicated in white font. Scale bar 25μm.

**Figure 6.**
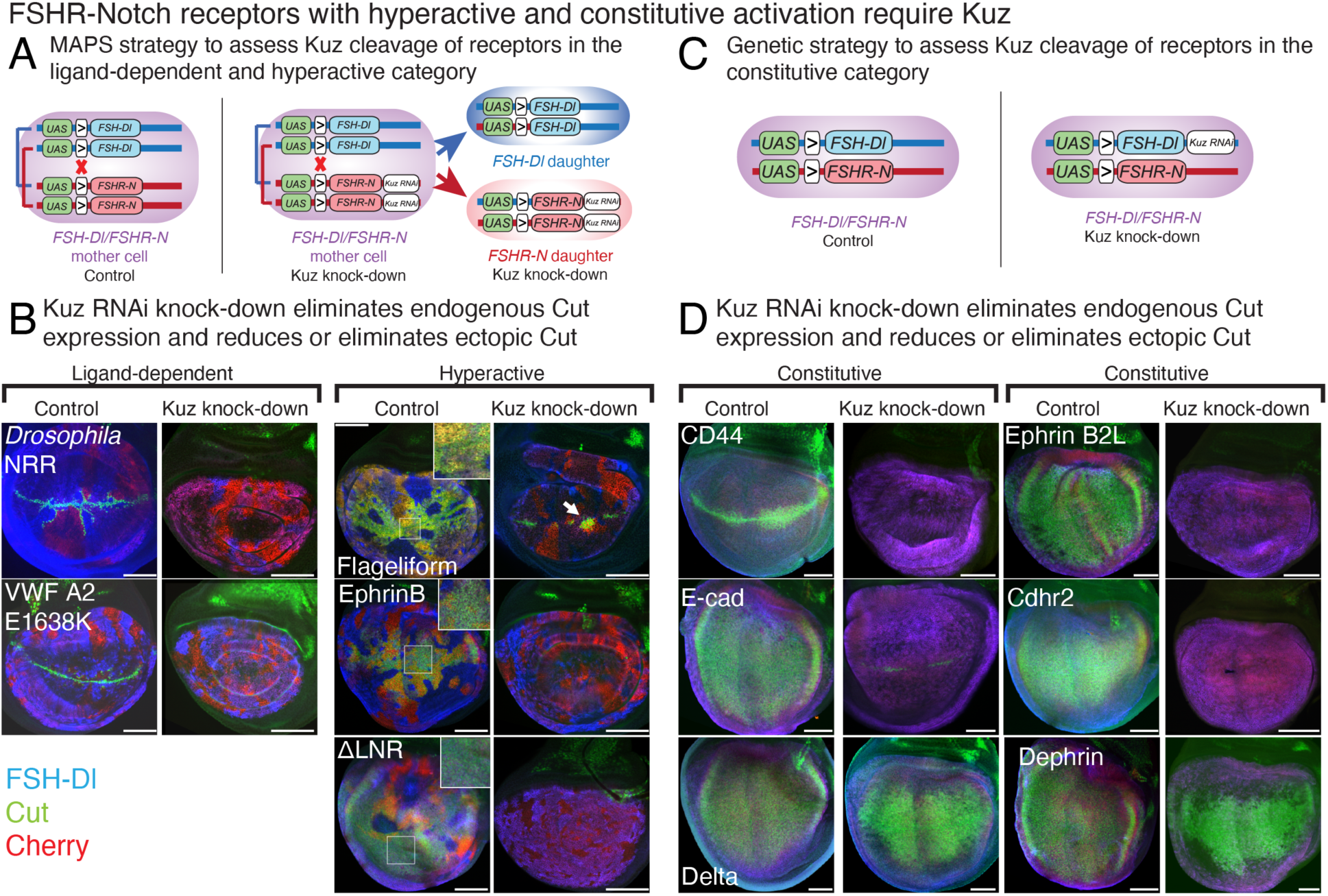
FSHR-Notch receptors with hyperactive and constitutive activation require Kuzbanian. **A**. Mitotic recombination using MAPS to compare the consequences of knocking down Kuz activity in cells that express ligand-dependent versus hyper-sensitive FSHR-Notch receptors. Left: Control genetic scheme using the ‘twin spot’ MAPS paradigm, with a *UAS>FSHR-Notch/UAS>FSH-Dl* parental cell and twin spot daughter cells homozygous for either *UAS>FSHR-Notch* or *UAS>FSH-Dl*. Note that in transheterozygous (mother) cells, canonical FSH-Dl and FSHR-Notch proteins mutually “cis-inactivate” each other. As a consequence, Cut activation is only induced at mosaic interfaces in which one or both clonal cell populations are homozygous for either *UAS>FSHR-Notch* or *UAS>FSH-Dl*. Right: Experimental genetic scheme to generate FSH-Dl/FSHR-Notch interfaces in which Kuz is knocked down in all FSHR-Notch expressing cells. This genotype is identical to the control scheme, except for the addition of *UAS.Kuz-RNAi* distal to *UAS>FSHR-Notch*. **B**. For ligand-dependent benchmark receptors (carrying the native Notch NRR or the A2^E1638K^ switch domains) in control discs in which Kuz is uniformly active (left most column), ectopic Cut is produced at mosaic borders at which one or both clonal cell populations are homozygous for either *UAS>FSHR-Notch* or *UAS>FSH-Dl* (first column). In the Kuz knock-down discs (second column), native DSL/Notch signaling is sufficiently impaired that Cut is not induced along the DV boundary, and FSH-Dl/FSHR-Notch signaling also fails to induce Cut for the Notch NRR or A2^E1638K^ receptors. For the hyperactive receptors carrying the Flageliform, EphrinB and ΔLNR candidate domains, control wing discs show extensive Cut activation in most FSHR-Notch cells (insets), but not in homozygous *UAS>FSH-Dl* clonal populations, which lack the *UAS>FSHR-Notch* transgene (third column). We infer that this Cut activation reflects constitutive activity of the receptor. In contrast, the experimental Kuz knock-down discs do not express Cut, except in homozygous *UAS>FSHR-Notch* clones encoding the Flageliform receptor positioned close to the DV border (arrow), which we interpret as due to residual Kuz activity. **C**. Genotypes for testing if Kuz is required for the transducing activity of constitutively active FSHR-Notch receptors. Both control (wild type for *Kuz*) and experimental (Kuz knock-down) animals are *UAS>FSHR-Notch/UAS>FSH-Dl*, with experimental animals also carrying *UAS.Kuz-RNAi* in *cis* with *UAS>FSH-Dl*. D. Cut is uniformly expressed throughout most or all of the wing pouch of control animals, except in the case of the CD44 FSHR-Notch receptor, where Cut is ectopically expressed only in the vicinity of the DV border (from which we infer that it has weak constitutive activity, sufficient only to boost the higher levels of native DSL/Notch signaling in the vicinity of the border to generate an ectopic Cut response). In experimental animals with Kuz knocked-down, the Cut response is abolished for the CD44, E-cad, Ephrin B2L and Cdhr2 FSHR-Notch receptors, but not the Delta and Dephrin receptors. Scale bar 50μm.

For 6 of the constitutively active receptors, we used a simpler, non-MAPS strategy (Fig. 6C). In this case, 4/6 of these receptors behaved as strictly Kuz dependent. However, the remaining 2/6 receptors, from *Drosophila* Delta and ephrin, behaved as if cleaved in a Kuz-independent manner (Fig. 6D).

Hence, except for two possible exceptions, activation of all of the chimeric FSHR-Notch receptors tested, whether ligand-dependent or constitutive, depends on the Kuz protease.

## Discussion/Conclusions

We present a readily scalable, *Drosophila* screen to identify protein domains that can substitute for the Negative Regulatory Region (NRR) of Notch and activate the receptor in response to Epsin-mediated ligand endocytosis. In an initial, proof-of-principle test of 43 candidate domains, around one quarter (11/43) have this capacity and comprise a surprisingly diverse repertoire of seemingly unrelated structural motifs. Further, in every case, activation requires Kuzbanian (Kuz), the sole *Drosophila* ADAM10 protease responsible for cleaving the NRR to activate the native receptor.

These results (i) validate the screen, (ii) provide insight into the structural basis and cleavage of force-sensitive proteolytic switches by ADAM10 proteases, (iii) provide the means to identify new proteolytic switches for synthetic Notch biomedical applications, and (iv) raise the possibility that force-sensitive signaling mechanisms may be more prevalent than currently appreciated.

### A scalable screen to identify protein domains that are cleaved in response to force

In our initial screen, we were able to quickly and unequivocally identify around half of the candidate ptoteolytic switches (23/43) as capable of mediating Notch activation when introduced in place of the native, force-dependent NRR domain. Of these, 11/23 exhibited ligand-dependence, whether stringent (8/11) or more relaxed (3/11), whilst the remaining 12/23 conferred constitutive activity. We found that designations made on the basis of alterations in adult wing morphology following a simple genetic cross, were validated by analysis of Notch target gene expression in wing imaginal discs.

Further, we were able to test and establish the requirements for Epsin-dependent ligand endocytosis and Kuz proteolysis for each of the 11 ligand-dependent proteolytic switches. Finally, we optimized the methods used to conduct the screen, providing the potential to scale up the approach to assess hundreds rather than tens of different candidate proteolytic switches.

### A large and diverse collection of sequences can behave as force-sensitive proteolytic switches

Our initial choice of candidate proteolytic switches was biased, in part, by prior evidence that other juxtacrine signaling systems aside from DSL/Notch might likewise depend on force-dependent cleavage mechanisms. These include Eph/Ephrin, Frazzled/ DCC and the Dscam family of proteins, all of which have roles in neuronal guidance (*38*). Some of these proteins share phenomenology with the Notch receptor, including γ-secretase cleavage following ADAM10 cleavage (*25*), transendocytosis of parts of the ligand/receptor bridge from one cell to another (*39, 40*), and intracellular domains that may act as transcription factors (*37, 41*). Other protein domains we tested were chosen based on their structural or functional properties [From (*28*); Flagelliform], or serendipitous observations (for example Venus fluorescent protein). However, regardless of the justifications for choosing the candidates, the domains that tested positively for ligand, Epsin and Kuz dependent activation exhibit remarkable structural diversity as assessed by AlphaFold predictions.

That we observe such diversity amongst proteolytic switches suggests that the sequence and structural space for Kuz-dependent force sensing domains is large. This could suggest that factors other than solely exposure of the substrate cleavage site are important in the precise regulation of cleavage in different contexts. Possibilities include the action of tetraspanin proteins that place the protease active site at the substrate in a position poised for cleavage (*42*) and autoinhibitory structures within the protease that potentially have a capacity to respond to mechanical tension (*43*). Curiously, the ADAMTS13 protease that cleaves native Von Willibrand factor docks with the substrate, poised just above the occluded cleavage site to execute the cleavage if the site is rendered accessible by torsional strain (*44*). Hence, at least some of the specificity for Kuz/ADAM10 cleavage of the Notch NRR may similarly depend on the docking of the protease to the receptor, whether directly or through a bridging protein. Since the proteolytic switches we have identified all operate in the context of the FSH-Dl/FSHR-Notch signaling paradigm in which the receptor retains only the transmembrane and intracellular domains of native Notch, any such docking of Kuz would presumably have to be to these domains.

Three other aspects of our findings are worthy of note.

First, the 11 candidate domains that tested positively for ligand, Epsin and Kuz dependence appear to fall on a spectrum reflecting requirements for different levels of Epsin-mediated, endocytic force. Specifically, 5/11 are strictly dependent on Epsin-mediated ligand endocytosis, as determined by their failure to mediate Notch activation in response to modified forms of the ligand that have limited or no ability to enter the Epsin pathway. In contrast, 3/6 of the remaining candidate domains can activate Notch, albeit weakly, in response to these same modified ligands, a result we interpret as evidence that these domains are tuned to a lower force threshold. Finally, the remaining three ligand-dependent domains appear to confer both a heightened response to ligands that have restricted access to the Epsin pathway as well as modest constitutive activity. Accordingly, we infer that they constitute a distinct class of hyperactive proteolytic switches tuned to an even lower level of force.

Second, the hyperactive class includes a partially deleted form of the Notch NRR that retains little else except the defined Kuz cleavage site, suggesting that the peptide sequence containing the cleavage site retains some residual capacity to function as a force-sensitive proteolytic switch. The hyperactive receptors also produce a response to FSH-Dl multiple cell diameters away from the clonal interface. We have recapitulated this phenomenon using simulations of the activation of especially potent receptors that incorporate cell proliferation and the displacement of receiving cells away from the ligand expressing cells (*45*). Alternatively, a long-distance response to FSH-Dl could reflect heightened sensitivity to signaling mediated by filopodial extensions that can extend several cell diameters away from signal-sending and/or signal-receiving cells, as documented for native DSL/Notch signaling (*46, 47*).

Third, the properties of all of the candidate switch domains contrast with properties of the native NRR from the *C. elegans* Notch protein LIN-12 (*31*). In this case, the receptor can be strongly activated by ligand, whether or not it has access to the Epsin pathway but appears devoid of activity in the absence of ligand. We speculate that the LIN-12 NRR comprises a special kind of proteolytic switch that has been evolutionarily selected to respond non-linearly to force, such that it can be activated even in the absence of Epsin-mediated endocytosis while remaining strongly ligand-dependent.

### Prevalence of force-sensitive intercellular signaling mechanisms

A major motivation for screening for new NRR-like proteolytic switches is the conjecture that force-dependent Kuz cleavage may apply to other juxtacrine signaling interactions in addition to DSL/Notch signaling. Of particular interest, four of the ligand-dependent switches identified in the screen (Dscam, DCC, EphB2, DEph) function in contact-dependent signaling between neurons, a context in which there is evidence that pico-Newton forces can be generated across ligand/receptor intercellular bridges, whether by Epsin-mediated ligand endocytosis (*48*) or other means (*49*). A mechanosensitive cleavage mechanism in this context could have important functional consequences, such as the release of intracellular domains that act as transcription factors (*41, 50*) or the transendocytosis of the bridge from one cell to another (*40*). Overall, the success of this initial screen in identifying a diverse repertoire of switches is consistent with mechanosensitive ADAM cleavage being a general principle of juxtacrine communication. Indeed, one clear implication is that it will be difficult, if not impossible, to anticipate the presence of such a switch in any given signaling protein by its sequence or structural properties. Instead, identification will depend on functional tests, ideally in vivo.

Increasing the repertoire of proteolytic switches for synthetic Notch biomedical applications.

Synthetic Notch (synNotch) technologies employing chimeric ligand and receptors to generate new, heterologous signaling systems offer significant potential for disease therapies and tissue engineering (*29, 30*). As the in vivo *Drosophila* screen presented here is both effective and scalable, it provides a new approach to identifying proteolytic switches that can be used to optimize and diversify the current repertoire of synNotch signaling systems, for example to generate synNotch receptors with distinct force thresholds or signal to noise ratios. As such it augments current approaches, for example using truncations of the NRR and heterologous cleavage domains in the context of the SNIPR synNotch receptor (*51*). As the screen is both effective and scalable, it should make it possible to identify new kinds of proteolytic switches, such as the LIN-12 NRR, to optimize and diversify the current repertoire of synNotch signaling systems. Finally, the use of a model organism to test synNotch components in vivo may help close the gap between the development of synNotch signaling systems and their clinical application (*52*).

## Methods

Further information and requests for resources and reagents should be directed to and will be fulfilled by the Lead Contact, Paul Langridge.

### Drosophila

All flies were maintained at 25°C on standard corn food. Fly stocks were maintained at 18°C, whilst experiments were performed at 25°C.

### *Drosophila* Transgenes

All ligand and receptor coding sequences, with the exception of FSHα, were inserted into a modified form of *pUAST-attB* (www.flyc31.org) that contains a single Flp Recombinase Target (FRT, ‘>’) positioned between the UAS promoter and the coding sequence, and the resulting *UAS>ligand* and *UAS>receptor* transgenes were introduced at a single genomic docking site, *attP-86Fb* located on the right arm of the third chromosome, oriented so that the promoter is centromere proximal to the coding sequence. A “no promoter” (*Ø*) element encoding the transcriptional terminating 3’ UTR of the *hsp70* gene is used in place of the UAS promoter in *Ø>ligand* transgenes (generated by Flp/FRT mitotic recombination in the male germline (*18*)). A single *UAS.FSHα,* transgene inserted by conventional P-element mediated transformation onto the X chromosome was used in all experiments.

The WT, KtoR, and ΔIC chimeric forms of Dl (Fig. 1A) are described in detail in (*18*). Briefly, the native extracellular domain of Dl was replaced in its entirety by the β subunit of human Follicle Stimulation Hormone (FSHβ) (*53*) and an HRP tag was inserted immediately downstream of the C-terminal YCSFGEMKE sequence of FSHβ and immediately upstream of the Dl transmembrane domain. FSHα was co-expressed with the FSHβ-Dl protein to reconstitute FSH, which is an FSHα/FSHβ heterodimer (for simplicity, we refer to all ligands as FSH-Dl chimeras and do not include the HRP tag in their designations).

The chimeric forms of Notch (Fig. 1A, B) are based on the FSHR-Notch receptor described in detail (*18*). Briefly the amino-terminal Epidermal Growth Factor (EGF) Repeat containing portion of the native extracellular domain of Notch was replaced by the ectodomain of FSHR. The extracellular domain was tagged by the insertion of Cherry just upstream of the juxtamembrane NRR domain.

For the screen, the native NRR was replaced with the coding sequence of the candidate domain of interest. In each case, the same restriction enzyme sites and junction sequences were used. The sequence of the NRR insertion position is between the upstream Cherry epitope tag and the downstream Notch TM domain are as follows: delyk…rppanvkyvit. The amino acid sequence of each of the ligand dependent and constitutive switch domains is given in table S3. The *UAS KuzHA* transgene and the *UAS KuzRNAi* originate from (*37*). Complete DNA sequences for both the ligands and receptors are available on request.

Analysis of signaling between dedicated ligand and receptor expressing cells.

Signaling between dedicated ligand and receptor cells was analyzed using Mosaic Analysis by Promoter Swap (MAPS; see (*18*) for detailed description). In essence, mitotic recombination across the FRTs in cells trans-heterozygous for *UAS>* and *Ø>* transgenes is induced in the presence of a *nub.Gal4* driver that acts in the developing wing. Upon the appropriate segregation of the recombined chromatid arms at the 4-strand stage, this generates mutually exclusive populations of ligand and receptor expressing cells; the ligand expressing cells are marked by staining for the HRP epitope tag (blue in all the Figures), and the receptor expressing cells are marked “black” by the absence ligand expression (and when needed, red, by the Cherry epitope tag on the receptor). Signaling is monitored by assaying for ectopic expression of Cut protein from the Notch target gene *cut* in receptor expressing cells that abut ligand expressing cells along the MAPS ligand/receptor mosaic interface.

To induce mosaics by promoter swap, first or second instar larvae of the appropriate genotype (*hsp70.flp UAS.FSHα/hsp70.flp* or *Y*; *nub.Gal4/+; Ø>*l*igand/UAS>*receptor or *UAS>ligand/Ø>receptor*) were heat shocked at 36 C for one hour and wing discs from mature third instar larvae were dissected, fixed, stained for Cut and HRP expression and mounted for confocal microscopy [commercially available Mouse monoclonal anti-Cut (Mab Developmental Studies Hybridoma bank 2B10, 1/100), and Rabbit anti-HRP (Abcam ab34885, 1/1000) antisera were used in combination with Alexa 488 and 633 conjugated labelled secondary antisera following standard protocols, for example as in (*18*)]. The Cherry epitope tag was also readily visualized in these preparations, although only shown in Fig. 5 and 6 (both the HA and HRP epitopes require rabbit monoclonal antisera for detection, precluding our use of the anti-HRP antisera to mark the “ligand” clones in this context). The intensity of the Cherry signal was similar for all forms of the receptors analyzed. Expression levels of the receptor were not further considered since we compared only relative levels of cleavage of the same receptor presented with different ligands. It is also possible that receptors scored refractory to cleavage were not strongly expressed.

### Quantification and Statistical Analysis

In all *Drosophila* MAPS experiments, most if not all the imaginal wing discs contained several mutually exclusive subpopulations of ligand and receptor expressing cells within each wing primordium. In all cases, the results of each ligand/receptor combination are presented by representative images as well as quantitative analysis. As noted in the main text, Cut expression is normally induced in “border” cells flanking the DV compartment boundary in response to native DSL/Notch signaling which peaks in the vicinity of the boundary and declines in a graded fashion as a function of distance from the boundary (*54*). Signaling between FSH-Dl and FSHR-Notch chimeras along MAPS mosaic borders located away from the DV boundary induces Cut when the combined inputs of the native and chimeric ligand/receptor interactions reach the levels normally required to induce Cut in DV border cells. Accordingly, the distances over which MAPS mosaic borders can induce Cut away from the boundary provide an indication of the strength of the signaling interaction between the chimeric ligand and receptor.

As depicted in Figure 3A, we measured and averaged the maximum distances from the DV boundary over which ectopic Cut was detected along 1-3 MAPS mosaic borders in each of at least 4 and typically many more discs for each genotype (power analysis>0.95, the numbers of discs analyzed for each ligand/receptor pair are presented in Fig. 3 and 4). As disc size varies at least in part because of extra growth induced by ectopic Notch activation, we normalized the average of the distances obtained in each disc by dividing it by the width of the DV compartment boundary within the prospective wing (the “Cut distance/DV width ratio”; Fig. 3A, y-axis). Images were coded and measured by multiple individuals to eliminate bias.

Quantitation of the resulting ratios for each ligand/receptor pair are presented in Figures 3 and 4 in the form of boxplots in which each point depicts the average of the ratios measured for a single disc, and the mean, median, and 25-75% interquartile ranges of the ratios are depicted by a crosses, a wide horizontal line, and the tops and bottoms of each box, respectively. Statistical comparisons of the results obtained for different ligand/receptor pairs were performed by Brown and Forsythe tests followed by Dunnett TW post-hoc analysis using Graphpad software. Assessment of Cut expression following Kuzbanian RNAi expression was replicated in at least three separate experiments for each receptor presented.

### Colabfold Structural Predictions

For the AlphaFold computational analysis of protein structures (*55*), the open-source software ColabFold (*56*) was employed (versions 1.3.0 and 1.5.2). The notebook “ColabFold: AlphaFold2 using MMseqs2” was used through the Google Collaboratory platform to run AlphaFold with the following parameters: use_templates = false, use_amber = true, msa_mode = “MMseqs2 (UniRef + Environmental)”, model_type = “AlphaFold2-ptm”, num_models = 5, model_order = [1, 2, 3, 4, 5], num_recycles = 6, rank_by = “plddt,” max_msa = null, pair_mode = “unpaired + paired”.

The amino acid sequences of the ligand-dependent domains of interest were entered to ColabFold for structure prediction using the specified parameters. No templates were used in the predictions. The AlphaFold algorithm generates five different structural models for each individual input. The models are ranked from best to worst based on the overall predicted Local Distance Difference Test (pLDDT), AlphaFold’s per-residue confidence metric. Along with the structural models, the Predicted Aligned Error (PAE) of each model was visualized on a matrix. This metric implies a measure of confidence in the relative positions of residue pairs. The best confidence model prediction was chosen in each case and displayed in Figure S2.

## Supplementary Materials list

Figure S1. Sequence alignment of 28 amino acid region N-terminal to the transmembrane domain for FSHR-Notch receptors in distinct categories.

Figure S2. Structural predictions of each of the candidate domains cleaved in response to ligand.

Figure S3. Genetic scheme for producing Kuz RNAi knock-down using MAPS.

Table S1 List of domains tested from the SNAPs cell culture assay

Table S2 List of all candidate domains.

Table S3 Amino acid sequences of cleaved domains.

## Acknowledgements

Thanks to Dominic Huntley, Olivia Mosley, Abby Bryant, Navya Pampatwar, Dat Dang and Corey Andrews for technical assistance. **Funding:** Research reported in this publication was supported by the National Institutes of Health through the Institute of General Medicine of the under award number R35 GM127141 (to G.S) and the Institute of Child Health and Human Development under award number R21 HD107414 (to P.D.L.), in addition to support from the Roy and Diana Vagelos Precision Medicine Award (G.S., P.D.L.) and the Research Scholarship & Creativity Activity Program at Augusta University (to P.D.L.). The content is solely the responsibility of the authors and does not necessarily represent the official views of the National Institutes of Health. **Author Contributions:** Conceptualization, P.D.L. and G.S.; Investigation P.D.L, F.C.B., T.J., B.W., A.A., R.F.A., S.B.; Writing – original draft, P.D.L. and F.C.B.; writing – review and editing, P.D.L., F.C.B. and G.S., supervision, P.D.L., funding acquisition, G.S. and P.D.L. **Declaration of interests:** The authors declare no competing interests. **Data and Materials Availability:** All data needed to evaluate the conclusions in the paper are present in the paper or the Supplementary Materials. Resources and reagents available on request.

