## Supplemental Material for "An in vivo screen for proteolytic switch domains that can mediate Notch activation by force"

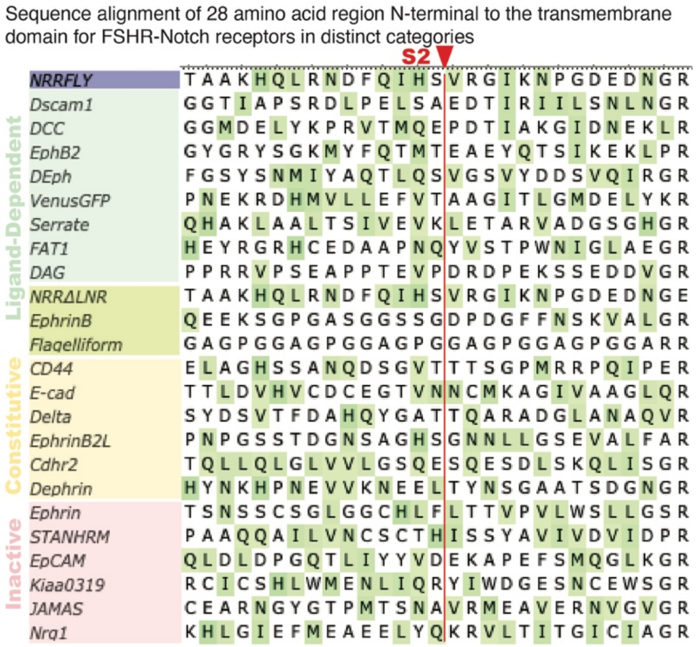


**Figure S1. Sequence alignment of 28 amino acid region N-terminal to the transmembrane domain for FSHR-Notch receptors in distinct categories.**

Alignment of 28 amino acids immediately N-terminal to a PPAN linker and the *Drosophila* Notch transmembrane domain, VKYVIT..... Hydrophobic amino acids favored by ADAM10 at the P1' site (amino acid immediately C-Terminal to the cleavage position) are labelled green. There is little correlation in the presence or position of these amino acids and the domain categorization - inactive (red), constitutive (yellow), ligand dependent (light green), ligand-dependent hyperactive (chartreus). Indeed, ligand-dependent proteolytic switches include sequences with similar amino acids at or close to the P1' position (DEph, Serrate, EphB2) and amino acids with no hydrophobic amino acids (flagelliform). See Table 1, S2 and S3 for more information about the selected candidate domains.

#### Structural predictions of each of the candidate domains cleaved in response to ligand

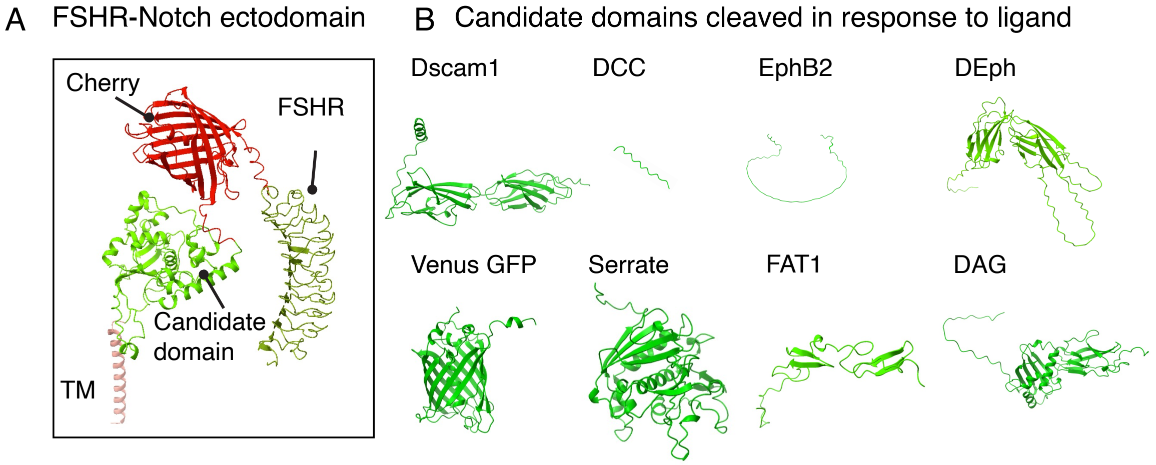


**Figure S2. Structural predictions of each of the candidate domains cleaved in response to ligand.**

**A.** High confidence structural predictions of domains showing ligand-dependent cleavage. The 3D structure of the entire extracellular region of canonical FSHR-Notch was predicted using AlphaFold [(*55*) see materials and methods].

**B.** This prediction was repeated for receptors with each of the candidate domains, and in each image only the candidate domain is shown and the remaining receptor hidden. Each candidate has a distinct structure with few commonalities. PDB files for each case are available.

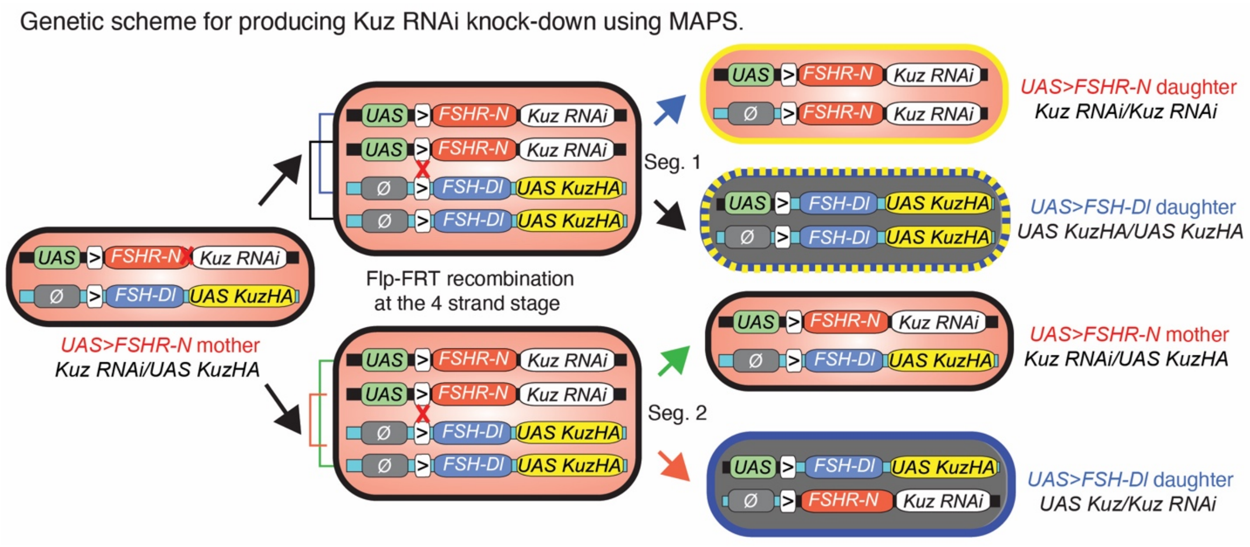


**Figure S3. Genetic scheme for producing Kuz RNAi knock-down using MAPS.**

Mitotic recombination using MAPS to remove detectable Kuz activity in sub-populations of FSHR-Notch cells. *UAS Kuz RNAi* was positioned distal to the *UAS>FSHR-Notch* transgene and *UAS KuzHA* distal to the *Ø>FSHβ-Dl* transgene. MAPS mitotic recombination at the four strand stage results in four possible segregant daughter cells (right and Fig. 5A), including one with the genotype identical to the mother cell.

List of domains tested from the SNAPs cell culture assay

| Name | Domain Source | Species | Description | Activity in SNAPS assay (*28*) | Activity category in the wing imaginal disc |
|---|---|---|---|---|---|
| FAT1 | FAT1 | Human | Atypical cadherin | Switch-like | Ligand-dependent class I |
| DAG | Dystroglycan | Human | Cell and ECM adhesion protein | Switch-like | Ligand-dependent class II |
| $A2^{E1638K}$ | Von Willibrand Factor | Human | Disease mutation of A2 domain | No activation | Ligand-dependent (*18*) |
| Cdhr2 | Cadherin-related family member 2 | Human | Cell adhesion protein | Switch-like | Constitutive |
| EpCAM | Epithelial Cellular Adhesion Molecule | Human | Adhesion molecule domain tested in SNAPS assay with no detectable cleavage. | No activation | No activation |
| KIAA0319 | Dyslexia-associated protein KIAA0319 | Human | Neuronal guidance protein domain tested in SNAPS assay with no detectable cleavage. | No activation | No activation |
| Muc13 | Mucin 13 | Human | Epithelial glycoprotein protein domain tested in SNAPS assay with no detectable cleavage. | No activation | No activation |

**Table S1.List of domains tested from the SNAPs cell culture assay.**

### List of all candidate domains

| | Name | Domain Source | Species | Size (#AA) | Description |
|---|---|---|---|---|---|
| Ligand-dependent | DCC | Deleted in Colorectal Cancer/ Frazzled | *Drosophila* | 18 | Neuronal guidance receptor |
| | EphB2 | Ephrin-type receptor B2 | Mouse | 110 | Neuronal guidance receptor |
| | DEph | Ephrin-type receptor | *Drosophila* | 250 | Neuronal guidance receptor |
| | Serrate | Serrate | *Drosophila* | 264 | Notch ligand |
| | FAT1 | FAT1 | Human | 93 | Atypical cadherin |
| | Dscam1 | Down Syndrome Cell Adhesion Molecule 1 | *Drosophila* | 220 | Neuronal guidance receptor |
| | Venus GFP | Venus GFP | *Aequorea victoria* | 242 | Entire coding region of Venus GFP |
| | DAG | Dystroglycan | Human | 261 | Cell adhesion and ECM binding protein |
| | $NRR^{\Delta LNR}$ | Notch variant | *Drosophila* | 45 | Variant of the NRR of Notch |
| | ephrinB | Ephrin B1 ligand | Human | 74 | Neuronal guidance protein |
| | Flagelliform | Flagelliform F40 | Spider | 44 | Spider silk – 40 PGGAG repeats |
| Constitutive | Dephrin | Ephrin B ligand | *Drosophila* | 211 | Neuronal guidance protein |
| | Robo | Roundabout 1 | *Drosophila* | 371 | Neuronal guidance receptor |
| | Nrg1 | Neuregulin (Nrg1 type III) | Human | 101 | Neuronal guidance protein |
| | Neogenin | Neogenin | Mouse | 367 | Neuronal guidance receptor |
| | Eprhin B2L | Ephrin B2 (Long) | Mouse | 342 | Neuronal guidance protein, juxtamembrane region and 2 fibronectin domains |
| | Ephrin B2S | Ephrin B2 (short) | Mouse | 110 | Neuronal guidance protein, juxtamembrane region |
| | EphB1 | Ephrin B1 receptor | Human | 124 | Neuronal guidance receptor |
| | Delta | Delta | *Drosophila* | 34 | Notch ligand |
| | APP | Amyloid-beta precursor protein | Human | 240 | Neuronal repair |
| | CD16 | CD16 | Human | 194 | Immune cell receptor |
| | CD44 | CD44 | Mouse | 461 | Cell adhesion glycoprotein |
| | Cdhr2 | Cadherin-related family member 2 | Human | 356 | Cell adhesion protein |
| Inactive | EphrinA2 | EphrinA2 Receptor | Mouse | 47 | Neuronal guidance receptor |
| | Fat | Fat | *Drosophila* | 211 | Atypical cadherin |
| | Filamin A20-21 | Filamin A 20-21 mechanosensitive region | Human | 370 | Actin crosslinking protein, Ig domain pair that acts as a mechanosensor domain requiring 5-10pN for unfolding |
| | Filamin GP1bα | Filamin A and glycoprotein protein (GP1bα) chimeric mechanosensitive region | Human | 293 | Actin crosslinking protein chimera, Ig domain pair that acts as a mechanosensor domain requiring 6-12pN for unfolding |
| | Nrg1 | Neuregulin Nrg1 (type III) | Mouse | 41 | Neuronal guidance protein |
| | STAN | Starry night (STAN) adhesion GPCR | *Drosophila* | 116 | Short sequence around ADAM proteolytic site of STAN |
| | PRP | Prion related protein (prp) | Mouse | 232 | Mouse prp |
| | STANHRM | Starry night (STAN) adhesion GPCR | *Drosophila* | 665 | Hormone receptor motif of STAN |
| | DSCAM | Down Syndrome Cell Adhesion Molecule 1 | Human | 232 | Neuronal adhesion and repulsion molecule |
| | Dachsous 1849-2057 | Dachsous isoform proteolytic cleavage region | *Drosophila* | 209 | Cadherin-related protein, isoform leading to N220/C210 processed form |
| | Dachsous 2323 - 2532 | Dachsous isoform proteolytic cleavage region | *Drosophila* | 210 | Cadherin-related protein, isoform leading to N270/C150 processed form |
| | EpCAM | Epithelial Cellular Adhesion Molecule | Human | 242 | Adhesion molecule domain |
| | SARS | SARS-Coronovirus-1 spike protein | SARS-COV-1 | 104 | SARS-COV1 subdomains 1 and 2 |
| | SARSRBD | SARS-Coronovirus-1 spike protein | SARS-COV-1 | 387 | SARS-COV1 subdomains 1 and 2 plus RGB domains |
| | COV2 | SARS-Coronovirus-2 spike protein | SARS-COV-2 | 108 | SARS-COV2 subdomains 1 and 2 |
| | COV2RBD | SARS-Coronovirus-2 spike protein | SARS-COV-2 | 392 | SARS-COV2 subdomains 1 and 2 plus RGB domains |
| | KIAA0319 | Dyslexia-associated protein KIAA0319 | Human | 236 | Neuronal guidance protein domain. |
| | Muc13 | Mucin 13 | Human | 249 | Epithelial glycoprotein protein domain. |
| | JAMAL | Junctional Adhesion Molecule A | Human | 206 | Cell adhesion molecule, including two Ig-like domains |
| | JAMAS | Junctional Adhesion Molecule A | Human | 107 | Cell adhesion molecule, including one Ig-like domains |

Table S2. List of all candidate domains.

Amino acid sequences of cleaved domains.

Ligand-dependent

Constitutive

| Name | Domain Source | Species | Amino Acid Sequence |
|---|---|---|---|
| Dscam1 | Down Syndrome Cell Adhesion Molecule 1 | *Drosophila* | VLAPPQSPHVTLSATTTDALTVKLKPHEGDTAPLHGYTLHYKPEFGEWETSEVSVDSQKHNIEGLLCGSR<br>YQVYATGFNNIGAGEASDILNTRTKGQKPKLPEKPRFIEVSSNSVSLHFKAWKDGGCPMSHFVVESKKRD<br>QIEWNQISNNVKPDNNYVVLDLEPATWYNLRITAHNSAGFTVAEYDFATLTVTGGTIAPSRDLPELSAEDTI<br>RIILSNLNG |
| DCC | Deleted in Colorectal Cancer/ Frazzled | *Drosophila* | VTMQEPDTIAKGIDNEKL |
| EphB2 | Ephrin-type receptor B2 | Mouse | APSAVSIMHQVSRTVDSITLSWSQPDQPNGVILDYELQYYEKQELSEYNATAIKSPTNTVTVQGLKAGAIY<br>VFQVRARTVAGYGRYSGKMYFQTMTEAEYQTSIKEKLP |
| DEph | Ephrin-type receptor | *Drosophila* | APTNLTLLFVDQTSAIISWSAPAKNESFSSETNSKIYHSDIVYKIKCNICSPNVVYNPSTDTFNETKITLTNLE<br>PVTTYTVQIHAINSVSHINEFKRHSNESSLVAVSDIVFSNTSLLNIPLDLNEVKTGQAEIVFTTESVLLSTVFN<br>LRILAITNKDADLEWDKPVQSDFPLEFYEVRWFPKVELDAINKSALNTKETKAHIVGLLENTEYGFQVRCK<br>TNNGFGSYSNMIYAQTLQSVGSVYDDSVQIRG |
| Venus GFP | Venus GFP | *Aequorea victoria* | MVSKGEELFTGVVPILVELDGDVNGHKFSVSGEGEGDATYGKLTLKLICTTGKLPVPWPTLVTTLGYGLQC<br>FARYPDHMKQHDFFKSAMPEGYVQERTIFFKDDGNYKTRAEVKFEGDTLVNRIELKGIDFKEDGNILGHKL<br>EYNYNSHNVYITADKQKNGIKANFKIRHNIEDGGVQLADHYQQNTPIGDGPVLLPDNHYLSYQSKLSKDPN<br>EKRDHMVLLEFVTAAGITLGMDELYK |
| Serrate | Serrate | *Drosophila* | DPKSACQNASNTISPYTALNRSQNWLDIALTGRTEDDENCNACVCENGTSRCTNLWCGLPNCYKVDPLS<br>KSSNLSGVCKQHEVCVPALSETCLSSPCNVRGDCRALEPSRRVAPPRLPAKSSCWPNQAVVNENCARLT<br>ILLALERVGKGASVEGLCSLVRVLLAAQLIKKPASTFGQDPGMLMVLCDLKTGTNDTVELTVSSSKLNDPQ<br>LPVAVGLLGELLSFRQLNGIQRRKELELQHAKLAALTSIVEVKLETARVADGSGHG |
| FAT1 | FAT1 | Human | LSPYCKDEPCKNGGTCFDSLDGAVCQCDSGFRGERCQSDIDECSGNPCLHGALCENTHGSYHCNCSHE<br>YRGRHCEDAAPNQYVSTPWNIGLAEG |
| DAG | Dystroglycan | Human | GGEPNQRPELKNHIDRVDAWVGTYFEVKIPSDTFYDHEDTTTDKLKLTLKLREQQLVGEKSWVQFNSNS<br>QLMYGLPDSSHVGKHEYFMHATDKGGLSAVDAFEIHVHRRPQGDRAPARFKAKFVGDPALVLNDIHKKIA<br>LVKKLAFAFGDRNCSTITLQNITRGSIVVEWTNNTLPLEPCPKEQIAGLSRRIAEDDGKPRPAFSNALEPDF<br>KATSITVTGSGSCRHLQFIPVVPPRRVPSEAPPTEVPDRDPEKSSEDDVG |
| $NRR^{\Delta LNR}$ | Notch variant | *Drosophila* | DIYDANYPGWNGGSGSTAAKHQLRNDFQIHSVRGIKNPGDEDNGE |
| ephrinB | Ephrin B1 ligand | Human | DPNAVTPEQLTTSRPSKEADNTVKMATQAPGSRGSLGDSDGKHETVNQEEKSGPGASGGSSGDPDGFF<br>NSKVALG |
| Flagelliform | Flagelliform F40 | Spider | GPGGAGPGGAGPGGAGPGGAGPGGAGPGGAGPGGAGPGGAR |
| Dephrin | Ephrin B ligand | *Drosophila* | CAPEDNNKTTALSNSKSVTDTGGAINVNIANNDESHVNSHGNNIAIGTNIGINGGQIIGGPQSAGIPINPLSG<br>NNNINGIPTTINSNIDQFNRIPIQPNIIGNHVGTNAVGTGIVGGGGIILTPGHAHGNINMLQPGRGGINGAYP<br>GHHHIQTGIRINNVPTQHNYPSHKGNANSNINGNDDHHHYNKHPNEVVKNEELTYNSGAATSDGNG |
| Robo | Roundabout 1 | *Drosophila* | PPGTPKVLNVSRTSISLRWAKSQEKPGAVGPIIGYTVEYFSPDLQTGWIVAAQRVGDTQVTISGLTPGTSY<br>VFLVRAENTQGISVPSGLSNTIKTIEADFDAASANDLSGARTLLTGKSVELIDASAINASAVRLEWMLHVSA<br>DEKYVEGLRIHYKDASVPSAQYHSITVMDASAESFVVGNLKKYTKYEFFLTPFFETIEGQPSNSKTALTYE<br>DVPSAPPDNIQIGMYNQTAGWVRWTPPPSQHHNGNLYGYKIEVSAGNTMKVLANMTLNATTTSVLLNNL<br>TTGAVYSVRLNSFTKAGDGPYSKPISLFMDPTHHVHPPRAHPSGTHDGRHEGQDLTYHNNGNIPPGDIN<br>PTTHKKTTDYLSGP |
| Nrg1 | Neuregulin (Nrg1 type III) | Human | VESNEIITGMPASTEGAYVSSATSTSTTGTSHLVKCAEKEKTFCVNGGECFMVKDLSNPSYLCKCPNEFT<br>GDRCQNYVMASFYKHLGIEFMEAEELYQKR |
| Neogenin | Neogenin | Mouse | VPSSLHVRPLVTSIVVSWTPPENQNIVVRGYAIGYGIGSPHAQTIKVDYKQRYYTIENLDPSSHYVITLKAF<br>NNVGEGIPLYESAVTRPHTDTSEVDLFVINAPYTPVPDPTPMMPPVGVQASILSHDTIRITWADNSLPKHQ<br>KITDSRYYTVRWKTNIPANTKYKNANATTLSYLVTGLKPNTLYEFSVMVTKGRRSSTWSMTAHGATFELVP<br>TSPPKDVTVVSKEGKPRTIIVNWQPPSEANGKITGYIIYYSTDVNAEIHDWVIEPVVGNRLTHQIQELTLDTP<br>YYFKIQARNSKGMGPMSEAVQFRTPKADSSDKMPNDQALGSAGKGSRLPDLGSDYKPPMSGSNSPHG<br>SPTSPLDSNML |
| Eprhin B2L | Ephrin B2 | Mouse | MAMARSRRDSVWKYCWGLLMVLCRTAISRSIVLEPIYWNSSNSKFLPGQGLVLYPQIGDKLDIICPKVDSK<br>TVGQYEYYKVYMVDKDQADRCTIKKENTPLLNCARPDQDVKFTIKFQEFSPNLWGLEFQKNKDYYIISTS<br>NGSLEGLDNQEGGVCQTRAMKILMKVGQDASSAGSARNHGPTRRPELEAGTNGRSSTTSPFVKPNPGS<br>STDGNSAGHSGNNLLGSEVALFA |
| Ephrin B2S | Ephrin B2 | Mouse | GQDASSAGSARNHGPTRRPELEAGTNGRSSTTSPFVKPNPGSSTDGNSAGHSGNNLLGSEVALFA |
| EphB1 | Ephrin B1 receptor | Human | PFPPQHVSVNITTNQAAPSTVPIMHQVSATMRSITLSWPQPEQPNGIILDYEIRYYEKEHNEFNSSMARSQ<br>TNTARIDGLRPGMVYVVQVRARTVAGYGKFSGKMCFQTLTDDDYKSELREQLP |
| Delta | Delta | *Drosophila* | DEESYDSVTFDAHQYGATTQARADGLANAQV |
| APP | Amyloid-beta precursor protein | Human | MVSKGEEDNSDVWWGGADTDYADGSEDKVVEVAEEEEVAEVEEEEADDDEDDEDGDEVEEEAEEPYE<br>EATERTTSIATTTTTTTESVEEVVRVPTTAASTPDAVDKYLETPGDENEHAHFQKAKERLEAKHRERMSQ<br>VMREWEEAERQAKNLPKADKKAVIQHFQEKVESLEQEAANERQQLVETHMARVEAMLNDRRRLALENYI<br>TALQAVPPRPRHVFNMLKKYVRAEQKDRQHTLKHFEHVRMVDPKKAAQIRSQVMTHLRVIYERMNQSLS<br>LLYNVPAVAEEIQDEVDELLQKEQNYSDDVLANMISEPRISYGNDALMPSLTETKTTVELLPVNGEFSLDDL<br>QPWHSFGADSVPANTENEVEPVDARPAADRGLTTRPGSGLTNIKTEEISEVNLDAEFRHDSGYEVHHQK<br>LVFFAEDVGSNKGR |
| CD16 | CD16 | Human | GMRTEDLPKAVVFLEPQWYRVLEKDSVTLKCQGAYSPEDNSTQWFHNESLISSQASSYFIDAATVDDSG<br>EYRCQTNLSTLSDPVQLEVHIGWLLLQAPRWVFKEEDPIHLRCHSWKNTALHKVTYLQNGKGRKYFHHN<br>SDFYIPKATLKDSGSYFCRGLFGSKNVSSETVNITITQGLAVSTISSFFPPGYQV |
| CD44 | CD44 | Mouse | HSKSHAAAQKQNNWIWSWFGNSQSTTQTQEPTTSATTALMTTPETPPKRQEAQNWFSWLFQPSESKS<br>HLHTTTKMPGTESNTNPTGWEPNEENEDETDKYPSFSGSGIDDDEDFISSTIASTPRVSARTEDNQDWT<br>QWKPNHSNPEVLLQTTTRMADIDRISTSAHGENWTPEPQPPFNNHEYQDEEETPHATSTTPNSTAEAAA<br>TQQETWFQNGWQGKNPPTPSEDSHVTEGTTASAHNNHPSQRITTQSQEDVSWTDFFDPISHPMGQGH<br>QTESKDTDSSHSTTLQPTAAPNTHLVEDLNRTGPLSVTTPQSHSQNFSTLHGEPEEDENHPTTSILPSST<br>KSGAKDARRGGSLPTDTTTSVEGYTFQYPDTMENGTLFPVTPAKTEVFGETEVTLATDSNVNVDGSLPG<br>DRDSSKDSRGSSRTVTHGSELAGHSSANQDSGVTTTSGPMRRPQIPE |
| Cdhr2 | Cadherin-related family member 2 | Human | VNDNPPTLDVASLRGIRVAENGSQHGQVAVVVASDVDTSAQLEIQLVNILCTKAGVDVGSLCWGWFSVAA<br>NGSVYINQSKAIDYEACDLVTLVVRACDLATDPGFQAYSNNGSLLITIEDVNDNAPYFLPENKTFVIIPELVL<br>PNREVASVRARDDDSGNNGVILFSILRVDFISKDGATIPFQGVFSIFTSSEADVFAGSIQPVTSLDSTLQGT<br>YQVTVQARDRPSLGPFLEATTTLNLFTVDQSYRSRLQFSTPKEEVGANRQAINAALTQATRTTVYIVDIQDI<br>DSAARARPHSYLDAYFVFPNGSALTLDE LSVMIRNDQDSLTQLLQLGLVVLG SQESQESDLSKQLISG |
| E-cad | Epithelial cadherin | Mouse | AEMDREDAEHVKNSTYVALIIATDDGSPIATGTGTLLLVLLDVNDNAPIPEPRNMQFCQRNPQPHIITILDPD<br>LPPNTSPFTAELTHGASVNWTIEYNDAAQESLILQPRKDLEIGEYKIHLKLADNQNKDQVTTLDVHVCDCE<br>GTVNNCMKAGIVAAGLQ |

**Table S3. Amino acid sequence of cleaved domains.**
